# Zeno Trap-enabled Quadrupole-Scanning Data-Independent Acquisition in nanoflow Quantitative Proteomics and Immunopeptidomics

**DOI:** 10.64898/2026.11.24.769014

**Authors:** Mohammadreza Dorvash, Ihor Batruch, Sri H. Ramarathinam, Nathan P. Croft, Patricia T. Illing, Jason Causon, Aaron Poth, Jarrod J. Sandow, Patrick Pribil, Katherine Tran, Stephen Tate, Anthony W. Purcell

## Abstract

We have investigated the quantitative and qualitative attributes of a scanning quadrupole data-independent acquisition (ZT Scan DIA) mode enabled on a SCIEX ZenoTOF 7600+ system using nanoflow chromatographic separations and targeted towards immunopeptidomic analysis. ZT Scan DIA incorporates increased ion utilization and MS/MS sensitivity introduced through the Zeno trapping modality, with quadrupole-resolved isolation windows in the data-independent acquisition mode. We demonstrate superior quantitative performance of ZT Scan DIA in sensitive nanoflow separations of tryptic peptides. Additionally, in LFQBench experiments, we demonstrate a remarkable quantitative accuracy and precision with experimentally measured precursor false-discovery rates below 1% using complex mixtures of yeast, human, and *E. coli*-derived tryptic digests when compared to conventional DIA experiments on the same platform. Having established its effectiveness in nanoflow proteomics analyses, we further investigated ZT Scan DIA for HLA-peptide characterization. These analyses highlighted the ability of the scanning quadrupole dimension to enrich the information needed for improved fragment-to-precursor assignment and quantitation of near-isobaric immunopeptides, should the search algorithms exploit it.

**Statement of Significance:** In this article, we evaluated Quadrupole-Scanning (ZT Scan) DIA on a ZenoTOF 7600+ platform, demonstrating its superior sensitivity, reproducibility, and quantitative accuracy in nano-flow proteomics analysis compared to ZenoSWATH DIA. This extended our earlier analysis in high throughput microflow applications to high resolution separations in nanoflow chromatography. We further highlighted using ZT Scan DIA for characterization and quantitative analysis of complex immunopeptidomics samples. We demonstrate that by incorporating quadrupole-scanning functionality, the Q1 dimension emerges from the sequential MS/MS scans in ZT Scan DIA, which can be further exploited to improve fragment-to-precursor alignment and fragment-ion prioritization for quantitation.

## INTRODUCTION

Immunopeptidomics provides a direct molecular readout of the peptides displayed by HLA molecules on the cell surface, capturing the antigenic landscape available for T-cell surveillance. This is particularly powerful in clinical samples, where genomics and transcriptomics can nominate potential antigens but cannot determine which peptides are actually processed, bound to HLA and presented to the immune system. By directly identifying naturally presented HLA class I and class II peptides, immunopeptidomics can reveal tumour-associated antigens, neoantigens, infection-derived peptides, autoimmune targets and non-canonical antigens arising from a variety of sources. Clinical immunopeptidomics remains technically demanding because the analytes of interest are scarce, diverse, and often need to be isolated and analysed from limited amounts of starting material (1). Unlike conventional proteomics, where abundant proteins dominate the signal, immunopeptidomics aims to identify short, low-abundance peptides that lack a common enzymatic cleavage pattern and are often unique to the HLA allotypes expressed by the tissue. This creates a challenging analytical problem in which peptide recovery, chromatographic efficiency, detection sensitivity, and statistical control and validity are all critical.

Nanoflow liquid chromatography coupled to high-resolution mass spectrometry is therefore essential for clinical immunopeptidomics. The improved ionisation efficiency of nanoflow electrospray provides substantially greater sensitivity than higher-flow-rate separations, which is particularly important when analysing scarce HLA peptides. Nanoflow separations also improve chromatographic resolution and sharpen the peaks, increasing the likelihood of detecting low-abundance immunopeptides against a complex background. While microflow approaches offer advantages in robustness, throughput and ease of operation, nanoflow remains difficult to replace when sample input is limiting and the primary objective is maximum antigen discovery depth.

For clinical translation, one must balance sensitivity with reproducibility, scalability and interpretability. Robust workflows require careful control of tissue processing, HLA immunoaffinity enrichment, peptide clean-up, LC-MS/MS acquisition, database construction and statistical validation. This is especially important when candidate antigens are intended to support vaccine design, T-cell monitoring, biomarker discovery or patient stratification. Advances in low-input sample preparation, high-resolution DIA, improved spectral libraries (1) and immunopeptidogenomic database strategies (2) are now making it increasingly feasible to apply immunopeptidomics directly to clinically relevant specimens, but nanoflow LC-MS remains central where antigen discovery from limited material is the primary goal. Converging with these requirements, the field of analytical biochemistry has been transformed by critical improvements in innovative data analysis strategies (3–5), and instrumentation and data acquisition modes (6, 7). These advancements, particularly in data acquisition and analysis, have enabled scientists to maintain technical validity and tackle complex biological questions (8). Since the introduction of the sequential window acquisition of all theoretical mass spectra (SWATH-MS) (9) as a universally applicable data-independent acquisition (DIA) mode, the field has seen several changes to enhance the sensitivity and specificity of peptide detection and quantitation. On one hand, innovations such as parallel-accumulation sequential elution and fragmentation (PASEF) were utilized to increase the precursor ion utilization by accumulating precursors at the instrument’s front-end, and to add the 4^th^ dimension of ion mobility to the LC-MS/MS data (10). ZenoTOF instruments benefit from the Zeno trap to maximize fragment ion utilization, to near 100% (11, 12). Enhanced ion utilisation has made it possible to design narrower isolation windows to exclude nearby analytes from interfering with the signal of the precursor of interest (13) without significantly compromising sensitivity. The ion processor of the Orbitrap Astral similarly accumulates ions in a low-pressure region of the ion trap, “crowd-controlling” them before releasing them into the TOF module (14, 15).

Analysis of DIA-MS data is principally founded on the targeted signal extraction, and either matching the spectra to peptide query parameters (PQPs) in a spectral library (i.e., peptide-centric approach) (9, 16), or generating “pseudo-DDA” spectra from correlating MS/MS signals, followed by a conventional database search (i.e., spectrum-centric approach) (16–18). The advancements in analyzing DIA data are twofold. Firstly, algorithms have evolved to deconvolve features across retention time, survey scan, product spectra, and ion mobility (if applicable) dimensions, and to confidently assign fragment ions to their precursor(s)-of-origin. An important concept that was adopted from targeted MS data analysis was the importance of fragment ion ratios to enhance confidence in identification. Thus, the second major revolution in DIA data analysis is machine-learning algorithms which emerged with the purpose of accurately predicting fragment ion ratios across a wide range of collision energies (19, 20). Eventually, the divide between the peptide- and spectrum-centric approaches has faded with the ever-improving accuracy of predicted fragment ratios. Now it is possible, particularly for tryptic peptides, to directly query the DIA data reliably using in-silico generated spectral libraries (21). Additionally, novel algorithms such as QuantUMS (5) were developed to prioritize precursor fragments for quantification, thereby maximizing the confidence of quantification across the analysis.

During the cycle time of conventional discrete window/stepping-DIA-MS, the quadrupole isolates each pre-determined m/z window for fragmentation and MS/MS acquisition, and moves up the m/z range one discrete window at a time. Isolated precursors in each window are subjected to the same collision energy, determined by the central m/z of each isolation window (9, 22). In scanning-quadrupole approaches like SONAR (23) and Scanning-SWATH (22), quadrupole parameters are adjusted in a continuous fashion, allowing the predefined isolation window to scan the precursor m/z range, without pausing or changing the isolation width. The collision energy is also smoothly ramped up for the higher m/z precursors. The rise and fall in the fragment intensities of each precursor in the product spectra provide an additional quadrupole-resolved dimension to this DIA acquisition strategy, commonly referred to as the Q1 dimension (6, 22). While this extra dimension makes the analysis of the scanning-DIA data much more complex, it enables better fragment-to-precursor alignment, which in turn increases the specificity/selectivity of the identification and quantification of each precursor. In ZT Scan DIA, Zeno trapping enhances fragment utilization, while the synchronous scanning of the quadrupole and time-of-flight pulsation enables alignment of the MS/MS signal with the quadrupole dimension, providing a more reliable precursor assignment (6). While several software packages can search the conventional discrete-window DIA, there are only a few software packages, such as DIA-NN (21) and PEAKS (versions 12 and above), that can currently utilize the Q1 dimension of ZT Scan DIA.

Here, we first confirmed the sensitivity and quantitative precision of ZT Scan DIA using different column loads of K562 tryptic digest separated with fast microflow liquid chromatography separations, and used Zeno SWATH DIA as the baseline. A similar detailed assessment of a multi-species mixture of human, yeast and E.coli tryptic digests in different ratios were used to assess quantitative precision across these proteomes in a LFQBench experimental format (24). These more nuanced experiments extend previously reported analysis (7), expanding upon the range of chromatographic gradients examined, the incorporation of experimentally derived spectral libraries for human, yeast, and *E. coli* to provide greater assignment confidence, and an extended ratiometric analysis of the LFQBench samples to further probe quantitative precision and accuracy under these chromatographic conditions. Such a detailed analysis further enabled more accurate comparison with new experiments designed to optimise nanoflow separations, which are ultimately better suited to immunopeptidomics, using both Zeno SWATH and ZT scan DIA modalities. Consistent with our earlier findings (7), ZT Scan DIA improved identification depth and quantitative performance in microflow workflows across an extended range of chromatographic conditions. Importantly, these benefits were also maintained under high-resolution nanoflow separations. Indeed, increasing chromatographic resolution amplified the performance advantages of both Zeno SWATH DIA and ZT Scan DIA, highlighting the complementarity between advanced acquisition strategies and improved peptide separation. Likewise, under nanoflow conditions improvements in quantitative accuracy and precision were seen using the multispecies LFQBench approach. Finally, we show that quantitative immunopeptidomics benefits substantially from ZT Scan DIA, particularly through improved fragment-to-precursor assignment enabled by the additional Q1 dimension, which helps resolve co-fragmented peptide signals that are often seen in complex HLA peptide mixtures.

## Materials and Methods

### Experimental design and statistical rationale

#### Assessment of Identification depth, quantitative reproducibility, and data completeness across different chromatographic conditions

We acquired Zeno SWATH and ZT Scan DIA data of tryptic digest of a human cell line (K562) proteome using a range of separation and acquisition parameters. The data were acquired using both separations under microflow conditions and a high-efficiency nanoflow separation in three technical replicates for each method combination. It should be noted that we excluded the use of ZenoSWATH DIA with a 13 ms accumulation time for the very short microflow separations (1- and 3-min gradients), as the combination of sharp chromatographic peaks and longer MS cycle times would have resulted in an insufficient number of data points per peak. We evaluated the identification depth at peptide and protein group level, as well as estimated the quantitative precision via calculating the coefficient of variation (CV%). Additionally, the data for one of the method combinations (5 ng loads separated using a 15 min gradient) was acquired with ten technical replicates to evaluate the data completeness of Zeno SWATH and ZT Scan DIA at similar identification depths.

#### Assessment of quantitative accuracy using a multi-species LFQBench design across different chromatographic conditions

To estimate the quantitative accuracy at the global level, we employed an LFQBench approach, using tryptic peptides of human, yeast, and *E. coli* mixed at known ratios; an index (or reference) mixture (HYE i0), three test mixtures with positive log_2_ fold changes (log_2_FC) for E. coli (HYE i+1, i+2, i+3), and three test mixtures with negative log_2_FC for E. coli (HYE i-1, i-2, i-3). The Zeno SWATH and ZT Scan DIA data were acquired in three replicates for each acquisition parameter combinations using both microflow and nanoflow separations. In addition, we fractionated the tryptic peptides of human, yeast, and *E. coli* separately using a high-pH kit and acquired the data in DDA mode to generate a spectral library. The LFQBench DIA data was searched using this high-pH spectral library. The log_2_FC of proteins and precursors for each species is calculated against the index group. Each of the six test groups is plotted separately, demonstrating a scatter plot and a box plot for the overall distribution of the measurements. The median of the measured log_2_FC values (M) and the log_2_-ratio of each species in comparison to the reference sample (R, dashed lines) are calculated for that group. The accuracy of quantitation is assessed by the closeness (Δ) of M and R values. An alternative way to assess quantitative accuracy is to evaluate the stoichiometry of the subunits of protein complexes. Here, we queried subunits of the constitutive 20S proteasome (only in yeast and human), for which seven α and seven β subunits are present at stoichiometric ratios.

#### Estimating the false discovery rates using the high-pH spectral library

The LFQBench DIA data were searched together with the K562-only files with similar acquisition parameters and separation conditions, using the high-pH spectral library described above. Since the spectral library was generated experimentally using separate fractions for each species, we were able to estimate the false discovery rate with more confidence than in previous comparisons (7). In addition, by searching the human-only and multi-species data together, we evaluated the FDRs with and without enabling the match-between runs option.

### Evaluating ZT Scan DIA for quantitative immunopeptidome analysis

We isolated class I immunopeptidome of the pancreatic cancer cell line Panc1. We acquired the LC-MS/MS data using Zeno SWATH and ZT Scan DIA to evaluate and compare their identification depth and quantitative accuracy and precision. Since the ZT Scan DIA methods had a restricted MS/MS isolation scan range, limited to 400 m/z to 900 m/z, we designed a variable-window ZenoSWATH DIA method based on Panc1 immunopeptide precursor ion m/z distribution, as described in detail below. Peptides were analyzed in three technical replicates for each DIA method. The data were searched using an experimentally acquired high-pH spectral library of Panc1 HLA-peptides (25).

#### Materials and reagents

LC-MS grade Water Optima® and Acetonitrile Optima® were purchased from Fisher Scientific. LC-MS grade Formic Acid (1 mL ampules), Trifluoroacetic Acid (TFA, 1 mL ampules), and High-pH Reversed-Phase peptide fractionation kit (#84868) were acquired from Thermo Fisher Scientific. Synthetic index retention time (iRT) peptides (26) were purchased from GL-Biochem (Shanghai). The LC-MS mobile phases A and B were prepared as 0.1% solutions of Formic Acid in water or Acetonitrile, respectively. An aqueous solution of 0.1% Formic Acid supplemented with 2% Acetonitrile was prepared (hereafter referred to as MS loading buffer) for use in peptide reconstitution and dilutions (unless otherwise specified). Commercially available tryptic digests of Human (Promega, K562 cell line digest, V6951), Yeast (Promega, Yeast digest, V7461), and *E. coli* (Waters MassPREP 186003196) proteome were purchased, reconstituted to a concentration of 1 µg/µL and separately aliquoted to 10 µg portions.

#### Serially diluted Human tryptic peptides

Aliquots (1 µg/µL) of tryptic digest of Human cell line (K562) proteome were further diluted to either 100 ng/µL or 125 ng/µL and were supplemented with iRT peptides at a 50 fmol/µL or 62.5 fmol/µL, respectively. The 100 ng/µL solution was serially diluted to 25, 2.5, and 0.5 ng/µL solutions. The 125 ng/µL solution was further serially diluted to 25, 12.5, 2.5, and 0.5 ng/µL. Peptides were immediately loaded into MS vials and queued for acquisition. For each run, 2 µL of the corresponding dilution was injected onto the columns.

#### Multispecies tryptic peptide mixtures for LFQBench analysis

LFQBench multispecies (Human + Yeast + *E. coli*) mixtures (hereafter referred to as HYE samples) were prepared as described in (24), with slight modifications. Seven mixtures (Fig. 3a) with arbitrary nomenclatures were prepared with predefined weight ratios of Human : Yeast : *E. coli*; i0 = 100:100:100; i-1 = 100:150:50; i+1 = 100:50:150; i-2 = 100:175:25; i+2 = 100:25:175; i-3 = 100:190:10; and i+3 = 100:10:190 (Fig. 3a). Briefly, the reference sample i0 has equal ratios of the three species. The next six samples contain an equal amount of Human peptides and varying amounts of Yeast and *E. coli*, with two-by-two reciprocal ratios. All the samples were prepared to a final concentration of 125 ng/µL supplemented with iRT peptides at 62.5 fmol/µL. An additional dilution of each HYE sample was prepared, with 25 ng/µL peptides and 12.5 fmol/µL iRT peptides. Peptides were immediately loaded into MS vials and queued for acquisition; for each run, 2 µL of the corresponding dilution was injected onto the columns. The equivalent amount of different injections of each sample is presented in Table S1.

#### High-pH Reversed-Phase (HPH-RP) fractionation of tryptic peptides

Human, Yeast, and *E. coli* tryptic peptides were fractionated separately using the HPH-RP stage columns, following the manufacturer’s instructions with slight modifications. Briefly, elution buffers were prepared by mixing the kit’s 0.1% Triethylamine (TEA) and Acetonitrile to final concentrations (v/v) of 10%, 14%, 18%, 22%, and 50% Acetonitrile. Three stage-columns were labeled, washed and conditioned with 0.1% TFA in Acetonitrile, and re-equilibrated with aqueous 0.1% TFA. 10 µg of each proteome was loaded separately to their corresponding column. After washing once with aqueous 0.1% TFA and once with pure water, peptides were sequentially eluted from the columns, collected, and stored. Prior to evaporative desiccation with a SpeedVac, samples were acidified with Formic Acid and frozen. Dry peptides were reconstituted with 12 µL of loading buffer and were spiked-in with 25 fmol/µL of iRT peptides. A total of fifteen library samples (five per species) were loaded to MS vials and queued for acquisition.

#### Isolation and preparation of HLA-peptides from Panc1

Pancreatic cancer cell line Panc1 was cultured using Dulbecco’s Modified Eagle Medium (DMEM) supplemented with additional necessary nutrients and chemicals (described in detail in (25)). Cells were maintained as adherent cultures at 37°C and 5% CO_2_. A pellet of 500 million cells was collected, followed by isolation of HLA-peptides according to procedures described in detail in (25, 27). HLA-peptide isolation was followed by desalting and cleanup, and evaporative desiccation. Dry peptides were reconstituted in 20 µL of MS loading buffer to achieve 25 million cell equivalents per µL (c.eq./µL). In addition, 2 µL of this peptide solution was taken and further diluted with MS loading buffer at a ratio of 1:10 to reach a 2.5 million c.eq./µL. Peptide solutions were spiked with iRT peptides at a 25 fmol/µL concentration. During LC-MS/MS runs, 1 µL of each sample was injected for data acquisition.

### LC-MS/MS

#### Liquid chromatography

We separated the peptides on an AQUITY M-Class UHPLC (Waters). The chromatographic conditions, including total run duration, gradient length, gradient start and stop B%, flow rate, and the RP-UHPLC column employed, are summarized in Table S2. Briefly, for micro-flow experiments, we used a 150 mm x 300 µm Kinetex (Phenomenex) column, operated at a flow rate of 5 µL/min and 40°C. Peptides were separated in a gradient of 3% to 30% mobile phase B, over 1, 3, 5, or 15 minutes of active separation. For nano-flow analysis of tryptic peptides (K562 digest and LFQBench experiments), we used a 600 mm x 75 µm Aurora FRONTIER XS column (IonOpticks), operated at a flow rate of 150 nL/min and 60°C. Peptides were separated using a gradient of 1% to 34% mobile phase B, actively separated over a 92 minute gradient (for multi-species spectral library generation) or a 30 min gradient (for serial dilutions and LFQBench experiments). For the analysis of the Panc1 immunopeptidome, we used a 150 mm x 75 µm Aurora ELITE XS column (IonOpticks), operated at a flow rate of 150 nL/min and kept at 60°C. Peptides were separated in a linear 15-min gradient of 1% to 34% mobile phase B.

### Tandem mass spectrometry

We employed a ZenoTOF 7600+ to acquire LC-MS/MS data. Parameter sets of MS/MS acquisition, including MS/MS acquisition mode, TOF MS and MS/MS scan ranges, isolation schemes, accumulation or dwell times, and total cycle times are summarized in Table S3.

Briefly, DDA data acquired for HPH-RP fractionated peptides utilized a 150 ms TOF MS survey scan of 400 to 1500 m/z, followed by 20 ms TOF MS/MS scans encompassing fragment ions of 140 to 1800 m/z. The top 45 most abundant precursors with a minimum intensity of 100 counts per second (cps) and charge states of +2 to +5 were selected for fragmentation and MS/MS acquisition; precursors were dynamically excluded for 5 seconds after 1 occurrence. The total cycle time for DDA acquisition was calculated to be 1259 ms.

For DIA analysis of tryptic peptides, the instrument began with a 50 ms TOF MS scan from 400 to 900 m/z. The ZT Scan DIA methods settings were predetermined based on chromatographic peak widths, expressed as full width at half maximum (FWHM), and were locked. The first ZT Scan DIA method was suitable for FWHM less than 1 second, using a sliding 5 m/z isolation window, scanning at 750 m/z per second with a dwell time of 6.7 ms and a total cycle time of 782 ms. The second ZT Scan DIA method was suitable for FWHM less than 4.5 seconds, using a sliding 10 m/z isolation window, scanning at 750 m/z per second with a dwell time of 13.3 ms and a total cycle time of 789 ms. In both methods, precursor isolation was restricted to between 400 and 900 m/z due to firmware constraints, and TOF MS/MS spectra were acquired between 140 and 1750 m/z. For ZenoSWATH DIA experiments, parameters were determined to be as comparable as possible to those used in ZT Scan DIA methods; 60 variable-width windows (width of 9 to 10 m/z) covering the 400 to 900 m/z range were used to isolate precursors for fragmentation and MS/MS acquisition. MS/MS accumulation time was set to 7 ms or 13 ms, yielding total cycle times of 786 ms or 1146 ms, respectively. TOF MS/MS spectra were acquired between 140 and 1800 m/z.

For DIA analysis of Panc1 HLA-bound peptides, the instrument began with a 50 ms TOF MS scan from 400 to 900 m/z. We employed the first ZT Scan DIA method (FWHM < 1s, Q1 window = 5 m/z, scan speed 750 m/z per second). We designed a similar ZenoSWATH DIA method based on Panc1 HLA-peptide precursors spanning 400 - 900 m/z, with 65 variable-width windows (minimum width of 5 m/z and 1 m/z overlaps). Precursors were isolated based on the isolation scheme in Table S4 for fragmentation, with an accumulation time of 7 ms and a total cycle time of 843 ms. TOF MS/MS spectra were acquired between 140 and 1800 m/z.

Throughout the manuscript, the term dwell time is used for both ZT Scan and ZenoSWATH precursor engagement times. Additionally, ZT Scan dwell times 6.7 and 13.3 ms are rounded to 7 and 13 ms, respectively, for aesthetic reasons in figures.

### Data processing and analysis

#### Spectral library generation

DDA .wiff files were transformed to centroided mzML using msConvertGUI in ProteoWizard 3.0.22189.522363d; Vendor-provided peak-picking algorithms were employed. FragPipe 22.0 was utilized to search the DDA files; MSFragger 4.1 search engine (28–30), EasyPQP 0.1.50, Philosopher 5.1.1 (31), MSBooster 1.2.3 (32), and DIA-NN (21) 1.9 were used in FragPipe to generate the spectral library. Table S5 summarizes the search parameters.

FASTA database preparation. The *Homo sapiens (n=20420), Saccharomyces cerevisiae (n=6060), and E. coli (K12 strain) (n=4530)* reviewed UniProt protein sequences (SwissProt) were downloaded on 2024.07.30. Eleven iRT peptide sequences (26) and forty-five non-Human, non-Yeast, non-E.coli proteins from the common repository of adventitious proteins (i.e., cRAP (33)) were appended to the database as contaminants. For FragPipe 22.0 search and library generation, the FASTA database had a total of 31066 target sequences and an equal number of reverse decoy entries.

#### DIA-NN 1.9.2 library-based searches

The DIA .wiff files were first converted to .wiff.dia formats using the DIA-NN 1.9.2 (21) GUI. Runs of each experimental comparison were searched with the corresponding settings and spectral libraries. For Human proteome serial dilutions, the spectral library described in (34) was used; deep learning was deactivated, and other settings are summarized in Table S6.

For LFQBench analyses, first, the HYE HPH-RP spectral library (generated in FragPipe, described *vide supra*) was imported to DIA-NN 1.9.2. Using the settings in Table S7, a predicted or *in silico* library was generated; deep learning was activated to refine the library, and re-annotation was deactivated to preserve the fractions and original species in which the peptides were observed. Experimental .wiff.dia files were searched using the refined spectral library and settings outlined in Table S7. We searched Panc1 HLA-peptidome data using the Panc1-specific immunopeptide spectral library described in (25). First, an *in silico* predicted (i.e., deep-learning refined) version of FragPipe library was generated using DIA-NN 1.9.2. For ZT Scan DIA files, the command-line option “--scanning-swath” was used. The remaining parameters were set as described in (25), and adjusted, summarized in Table S8.

### DIA-NN 1.9.2 experimental false discovery rate (FDR) estimation

In order to experimentally calculate the false discovery rates, the seven HYE experimental groups of each LC-MS/MS regimen (21 files) were searched together with the equivalent (acquisition regimen and on-column load matched) Human-only triplicates. The search strategy for these analyses is similar to that listed in Table S7, except that a separate *in silico* library is generated dedicated to FDR calculations. In addition, to assess how the MBR process affects the FDR, the main search was repeated once with and once without selecting MBR. This approach (use of a multi-species experimental spectral library) is adapted from Demichev et al. (21) with minor changes, including searching the Human-only samples together with LFQBench runs. FDR values are calculated as:

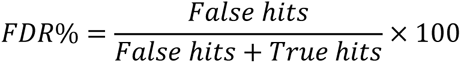

We calculated the protein-level FDR using the reported organism of the protein group determined during the protein inference. However, precursors were traced back to the files in which they were observed (by parsing the FragPipe psm.tsv), to ensure their species of origin. In other words, if a precursor is identified in a Human and Yeast protein FASTA sequence, but only traced back to Yeast’s HPH-RP fractions, it was only considered as a Yeast but not a Human peptide; if that precursor was traced back to both Human’s and Yeast’s fractions, it was considered as a homologous peptide. Both the falsely discovered and the homologous peptides were illustrated on the FDR plots. Of note, while this approach might underestimate the FDR due to the lower proportion of non-human precursors in the spectral library, the estimated values can be compared confidently across the search strategies.

### Precursors and proteins log-ratio diagrams and calculation of accuracy

Where required, chromatograms and spectra were inspected and extracted using the Explorer tab of SCIEX OS 3.4. An in-house Python script was used to curate the quantity matrices, perform calculations, and visualize the log-ratio plots. Coefficient of variation (CV%) was calculated as a measure of reproducibility (35, 36). In addition, to estimate the accuracy globally, the parameter Δ was defined (Fig. 3a) as the difference between the expected ratio of each species in each experimental group and the median of the measured quantities of that species’ peptides or proteins (24). We used Python 3.9 (in Anaconda environment) for data analysis and visualization; pandas 1.4.2, NumPy 1.21.5, matplotlib 3.5.1, and seaborn 0.11.2 were the libraries that we utilized via jupyterlab 3.3.2. Further visualizations were done in GraphPad Prism and Adobe Illustrator.

After searching the data files, log_2_-transformed ratios of precursors and proteins of each species in test samples (HYE i±3, i±2, and i±1) were calculated relative to the index sample i0. Log_2_-ratios of each entry are plotted on the y-axis against the log_2_-transformed intensity of that entry in i0 on the x-axis, in a scatter plot, as well as box-and-whisker plots depicting min, max, interquartiles, and the median of the log_2_-ratios of each species. In each test sample, the closeness (Δ) of the median of the measured log₂ ratios (M) for each species was calculated relative to its reference ratio (R); the R values are also visualized as dashed lines on the log-ratio plots. The accuracy of quantifications in each parameter set of ZenoSWATH and ZT Scan DIA can then be assessed by comparing the Δ values across the range of test samples and on-column loads (Fig. S4 and S7). This form of analysis gives insight into the overall systematic and random errors over the process of sample preparation, acquisition (LC-MS/MS), and quantification. In addition, to evaluate the quantitative accuracy based on the 20S proteasome subunit stoichiometry, the precursor and proteins of Human and Yeast subunits (Table S9) were illustrated with magenta stars in the log-ratio plots.

## Results

### ZT Scan DIA improves identification depth and quantitative reproducibility

Tryptic peptides of the human cell line K562 were prepared in serial dilutions to deliver on-column loads of 1, 5, 50, and 200 ng with a 2 µL injection. Samples were separated using a microflow (5 µL/min) configuration with linear gradients of 1, 3, 5, and 15 min duration. MS/MS data were acquired using discrete variable-window Zeno SWATH DIA or a continuous Q1-scanning ZT Scan DIA, with MS/MS accumulation or dwell times of 7 or 13 ms. Precursor (Fig. 1a) and protein group (Fig. 1b) identification counts across the evaluated parameter sets demonstrate that ZT Scan DIA achieved higher depths of identification, particularly during rapid chromatographic separations, with faster scan rates (i.e., shorter dwell times), and when input amounts were limited. Furthermore, when comparing identified protein groups and precursor counts, ZT Scan DIA consistently outperformed Zeno SWATH DIA with lower coefficients of variation (CV% less than 20% or 10% across replicates), demonstrating better quantitative reproducibility and precision (Fig. S1a,b). Furthermore, consensus between the two methods is seen at 44.5% to 75% of the peptide identifications, and 53.4% to 86.1% of the protein groups (Fig. S1c-d) as load is increased from 1 ng to 200 ng.

**Fig 1.**
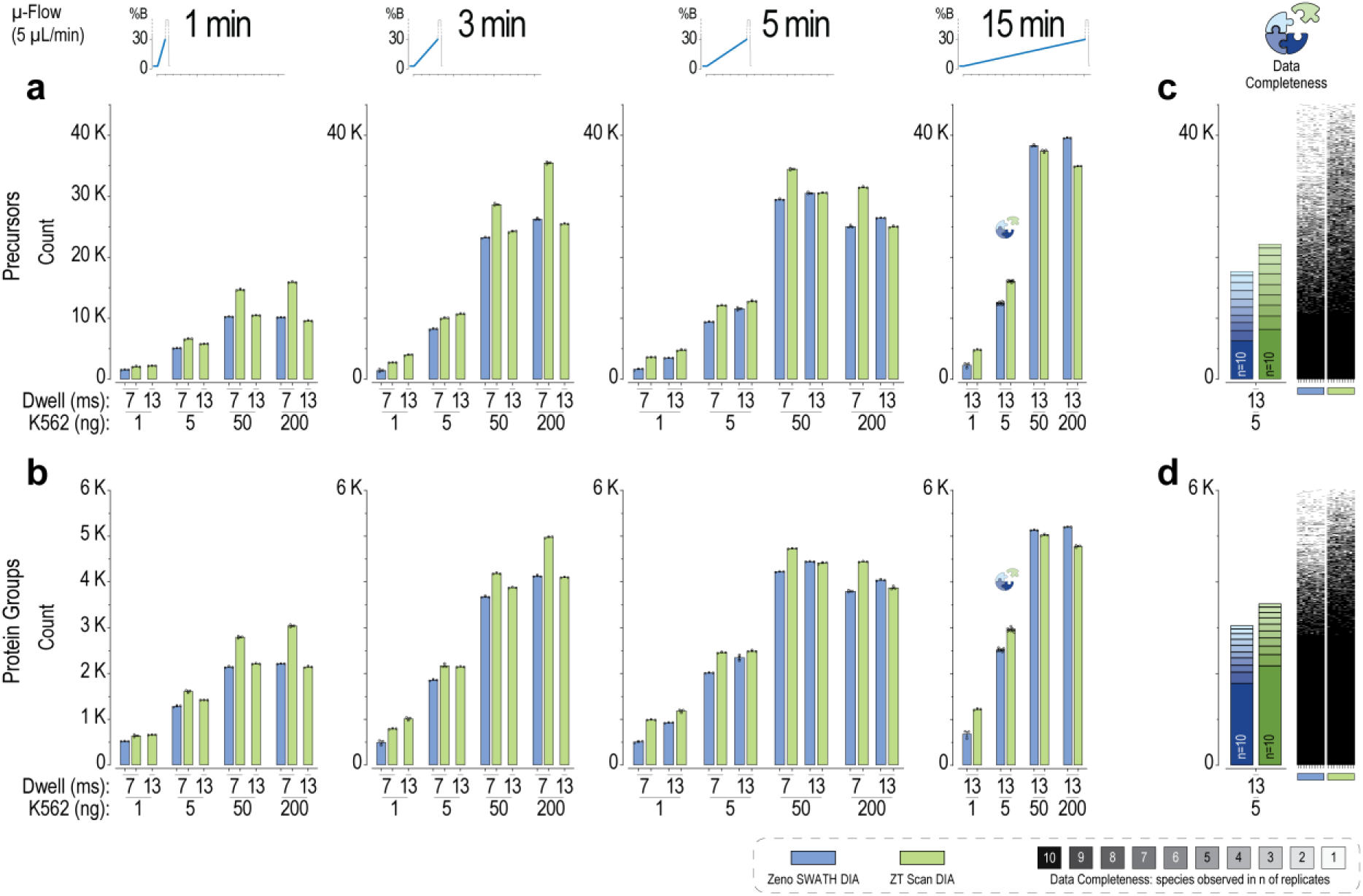
**ZT Scan DIA consistently improves the depth of identification** at **a**) precursor and **b**) protein group levels, with lower input amounts (on-column loads), shorter chromatographic gradients (i.e., higher throughputs), and shorter dwell times (i.e., faster scan speeds). Bar charts represent precursor and protein group counts of serially diluted (1, 4, 50, and 200 ng on column) Human K562 cell tryptic digest. Gradient lengths are indicated by the keys at the top of each precursor-protein plot pair (vertically); from left to right, 1, 3, 5, and 15 min gradient separations at a flow rate of 5 µL/min. The stacked bar charts and the binary heatmaps illustrate the data completeness at **c**) precursor and **d**) protein group levels are assessed by comparing ZT Scan DIA and Zeno SWATH DIA, both with 13 ms dwell times, and by acquiring 10 technical replicates of 5 ng loads with 15 min separations (the representative protein and precursor counts are also marked with the data completeness key). Each row of the heatmaps represents a precursor (**c**) or a protein group (**d**), each column represents a replicate, and black and white denote presence and absence, respectively. Note: Bar charts represent the mean of the measured values ± standard error of the mean (SEM) as error bars. Individual technical replicate measurements are depicted as points.

Comparing ZenoSWATH and ZT Scan DIA operated with the same gradient length, same sample loads, same dwell time (7 ms), and similar cycle times, both methods achieve similar estimates of FWHM and number of data points across the chromatographic peaks (Fig. S2a). When the longer dwell time (13 ms) of ZenoSWATH and ZT Scan DIA methods were compared at the same gradient lengths and loads, ZT Scan DIA acquired a greater number of points across the peaks (Fig. S2a). This was because, for the 13-ms dwell time, ZT Scan DIA offers a shorter cycle time compared to ZenoSWATH DIA. For the same reason, ZenoSWATH DIA overestimated the FWHM, as evidenced by deviations in the FWHM plots below the x=y line (Fig. S2a). Missing values are a common impediment to quantitative MS-based proteomics analyses (24, 37). Various technical and computational approaches, such as isobaric tag labeling and missing value imputation, have been developed to handle missing values in quantitative proteomics (37). DIA methods minimize the number of these missing values during data acquisition, which ultimately leads to a more robust LFQ analysis (38–40). Fig. 1c,d show that, for a given depth of identification, ZT Scan DIA achieves a higher data completeness at both precursor (Fig. 1c) and protein (Fig. 1d) levels compared to more traditional Zeno SWATH DIA acquisition. Finally, when comparing ZT Scan DIA with 7- and 13-ms dwell times across the same gradient lengths and sample loads, both methods yielded very similar cycle times, estimated comparable FWHM values, and acquired nearly the same number of data points per peak (Fig. S2b).

While microflow separation offers a higher throughput, it is almost exclusively surpassed by nanoflow setups in achieving chromatographic resolution and sensitivity (41). Thus, we decided to assess quantitative attributes of ZT Scan DIA acquisition using a high resolution Aurora FRONTIER XS (IonOpticks) UHPLC column operating at a flow rate of 150 nL/min and a 30 minute gradient. Serial dilutions of human K562 tryptic peptides demonstrated that ZT Scan DIA consistently identified more precursors (Fig. 2a) and proteins (Fig. 2b) across all on-column loads with more precise quantitation at both precursor and protein group levels as reflected by CV values below 10% and 20%, respectively. For instance, on average (expressed here as median and range), Zeno SWATH DIA and ZT Scan DIA identified 7155 (5974 to 7772) and 11641 (11326 to 12039) precursors, respectively, in only 1 ng of on-column material (Fig. 2a). Comparing to the same load separated for 5 min (microflow) with the same dwell times (7 ms), this is on average more than a 4-fold gain for Zeno SWATH DIA (7155 vs. 1660) and approximately 3-fold gain for ZT Scan DIA (11641 vs. 3627). In addition, since both DIA methods have similar cycle times, they also had similar FWHM values and acquired nearly the same number of data points across peaks for the different sample loads (Fig. S2c). It should be noted that in the smaller on-column loads (1 and 5 ng), the greater number of precursors detected by ZT Scan DIA (Fig. 2a) appeared as small deviations from the x=y line for both FWHM and acquired data points per peak (Fig. S2c).

**Fig 2.**
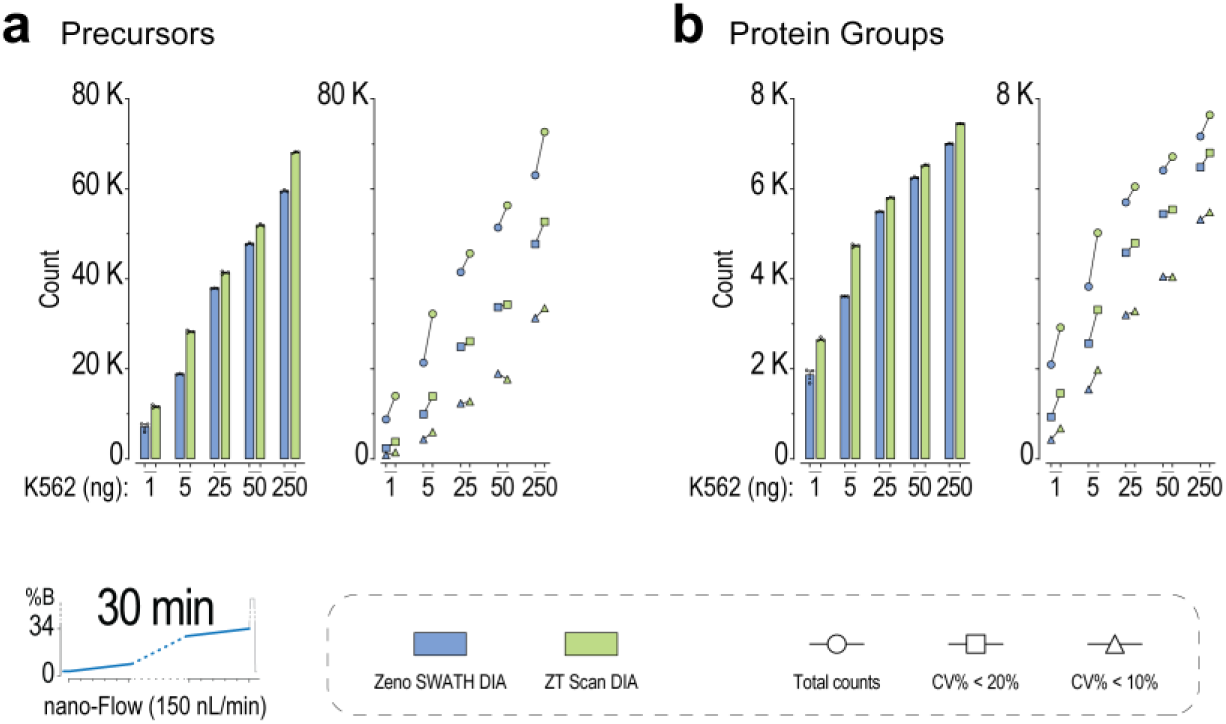
**ZT Scan DIA in a nanoflow separation configuration consistently increases the identification depth and quantitative precision of tryptic peptides**, demonstrated for **a**) precursors and **b**) protein groups. Note: Bars represent the mean of the measured values ± SEM. Separate technical replicate measurements are depicted as points, wherever possible. (Note for formatting: The figure best suits a width of ∼10.6 cm)

Furthermore, the utilization of longer chromatographic gradients, on average, resulted in higher precursor identification depth, which in turn, enabled higher overall protein sequence coverage (Fig. S3a). On the same note, the choice of ZT Scan DIA over Zeno SWATH DIA generally yielded higher sequence coverage, particularly with lower input samples (Fig. S3b); for instance, ZT Scan DIA achieves a median of 7 to 8-fold higher protein coverage for 1 ng of K562 across different gradient lengths and dwell times (Fig. S3b).

#### Generating a multispecies deep spectral library using high-pH off-line fractionation and nanoflow separation

To prepare a deep spectral library for LFQBench analyses (24), 5 high-pH reversed-phase (HPH-RP) fractions of cellular tryptic peptides of human, yeast, and *E. coli* origin were individually prepared. Data-dependent acquisition (DDA) was used to generate MS/MS spectra of peptides within these fractions using an extended 92-min gradient separation on an Aurora FRONTIER (60 cm x 75 μm) XS C18 UHPLC column (IonOpticks) at a flow rate of 150 nL/min. Each HPH-RP fraction yielded unique total ion chromatograms highlighting the orthogonality of the offline HPH-RP fractionation with this on-line separation modality (Fig. 3a). The data for all fractions were searched in FragPipe to generate a combined spectral library. Fig. 3b demonstrates the precursor counts per species for each HPH-RP fraction. Fig. 3c further elucidates the peptides that are unique to each species or are found across fractions of more than one species, i.e., are homologous. This species-resolved spectral library was generated to enable us to more accurately estimate the false discovery rates (FDR) independent of any protein inference procedure (21) (further details in Methods).

**Fig 3.**
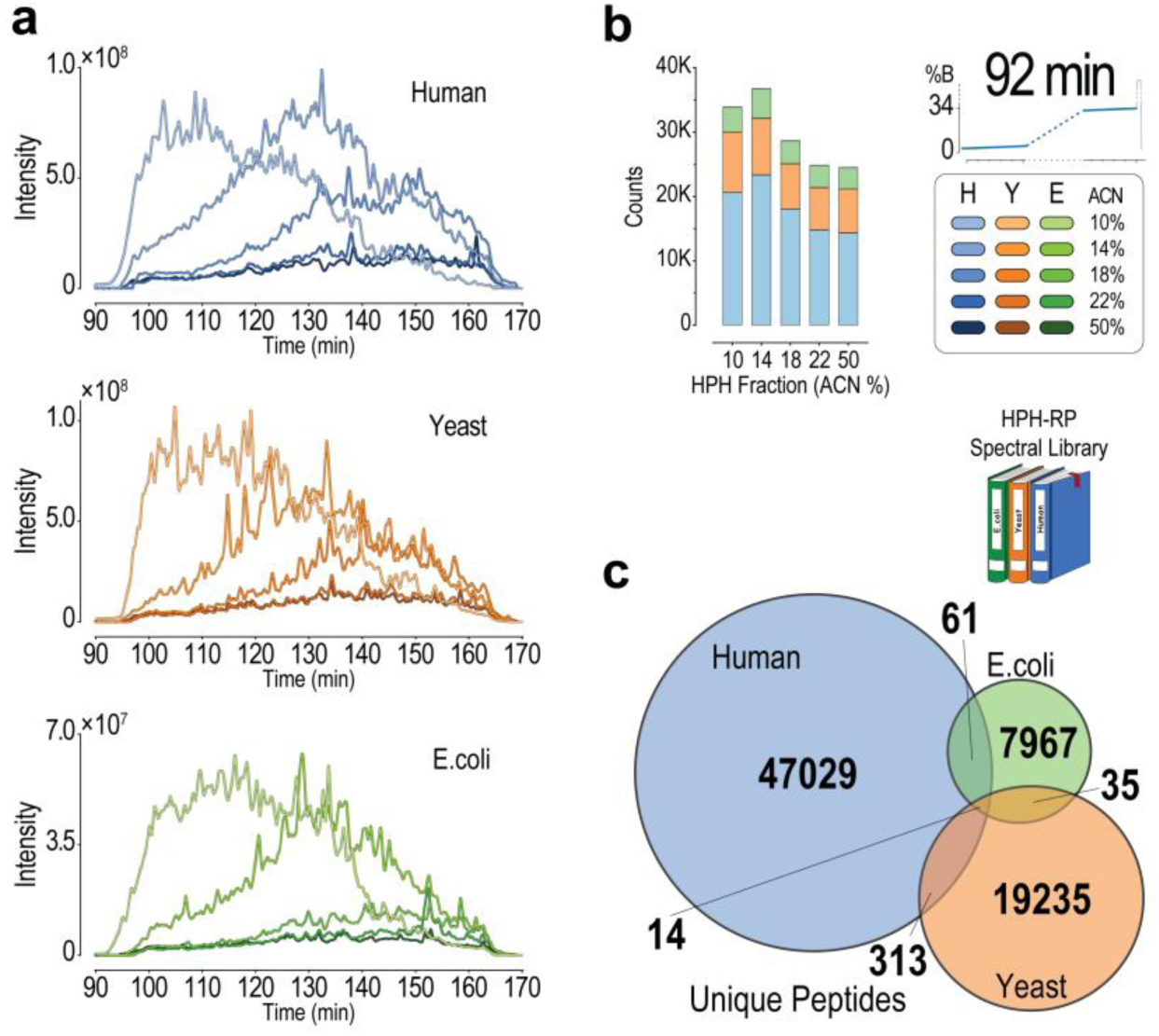
Generating and characterizing a multispecies deep HPH-RP spectral library to accompany LFQBench experiments. **a**) Tryptic digests of the three phylogenetically distant species (human, yeast, and E. coli) were separately fractionated using off-line HPH fractionation. Each line represents the TIC of a distinct HPH-RP fraction (equilibration times were removed from x-axes). As shown in the key, HPH-RP fractions are represented via shades of each colour. The distribution of identified peptides across HPH fractions (**b**) and the total number of peptides identified for each organism (**c**) are also presented. (Note for formatting: The figure best suits a width of ∼10.6 cm)

#### Nanoflow ZT Scan DIA provides robust quantitative performance across chromatographic gradients and sample complexities

Seven different predefined mixtures of human, yeast, and *E. coli* tryptic digests were prepared as per an extensive LFQBench protocol (24) (Fig. 4a,b). DIA LC-MS/MS data of each sample were acquired by injecting 250 ng of each mixed sample in three technical replicates, 30-min gradient separations (150 nL/min), and with 7 ms dwell times. Three technical replicates of on-column loads of 25 ng for the i0, i-1, and i+1 mixtures were also analyzed using the same LC-MS/MS methods.

**Fig 4.**
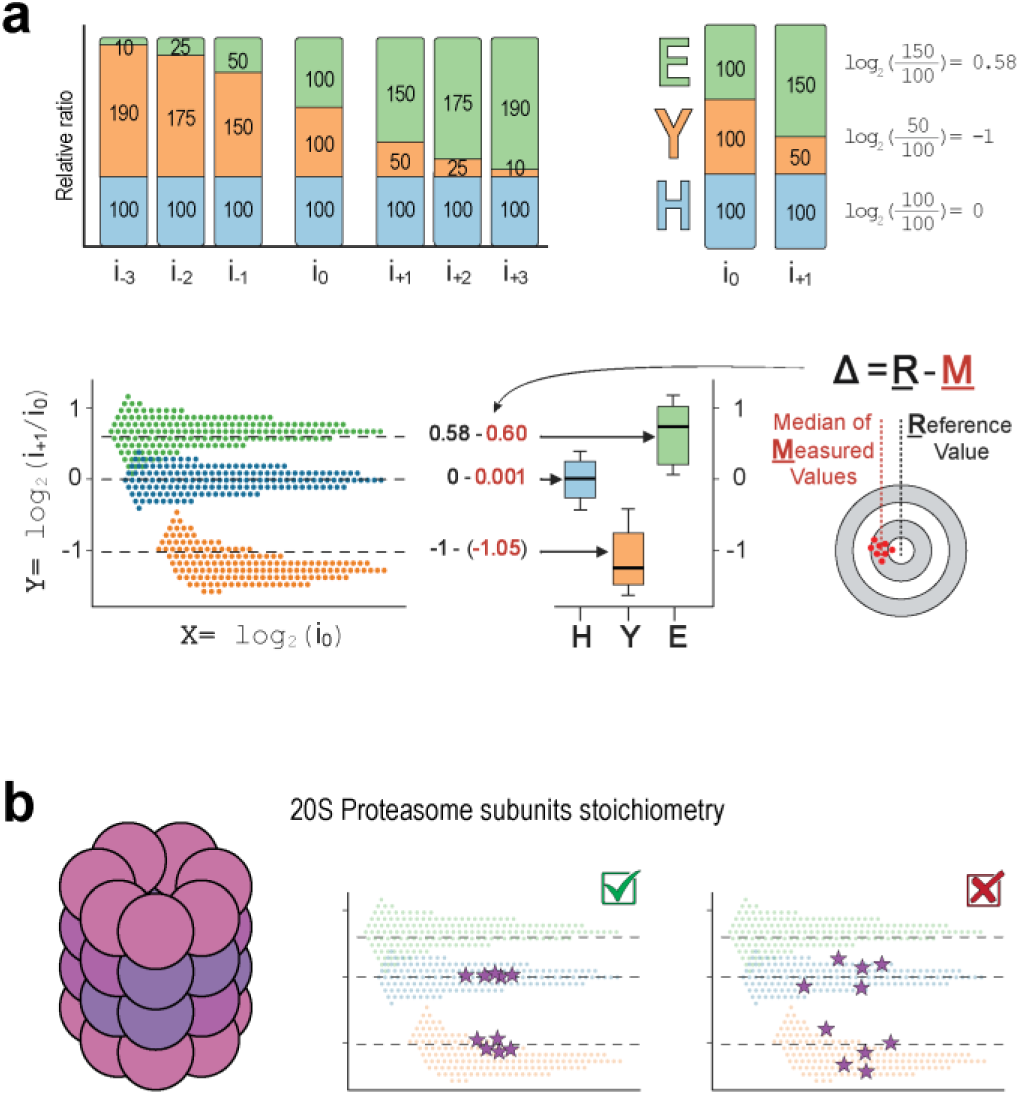
A brief schematic description of the LFQBench experimental design, with the approach to estimate quantitative accuracy at **a**) the global and **b**) in an alternative approach using the stoichiometry of subunits of human and yeast constitutive 20S proteasome subunits. (Blue, orange, and green circles are human, yeast, and E. coli precursors or proteins. Magenta stars represent 20S proteasome subunit precursors or proteins.) (Note for formatting: The figure best suits a width of ∼8.8 cm)

Fig. S4a,b show the counts of precursors and proteins identified, for each species, using high-efficiency nanoflow (30 min separation, 25 vs. 250 ng load, and 7 ms dwell time) setups.

The LFQBench ratio plots of nanoflow separations (Fig. 5) demonstrate that with the choice of a high-efficiency nanoflow chromatography, both Zeno SWATH DIA and ZT Scan DIA methods can achieve highly accurate quantifications . Nonetheless, ZT Scan DIA demonstrated greater overall accuracy, particularly for species present at the lower ratio. In the same fashion, the stoichiometric ratio of 20S proteasome subunits is measured with higher accuracy using ZT Scan DIA (Fig. 5, stars).

**Fig 5.**
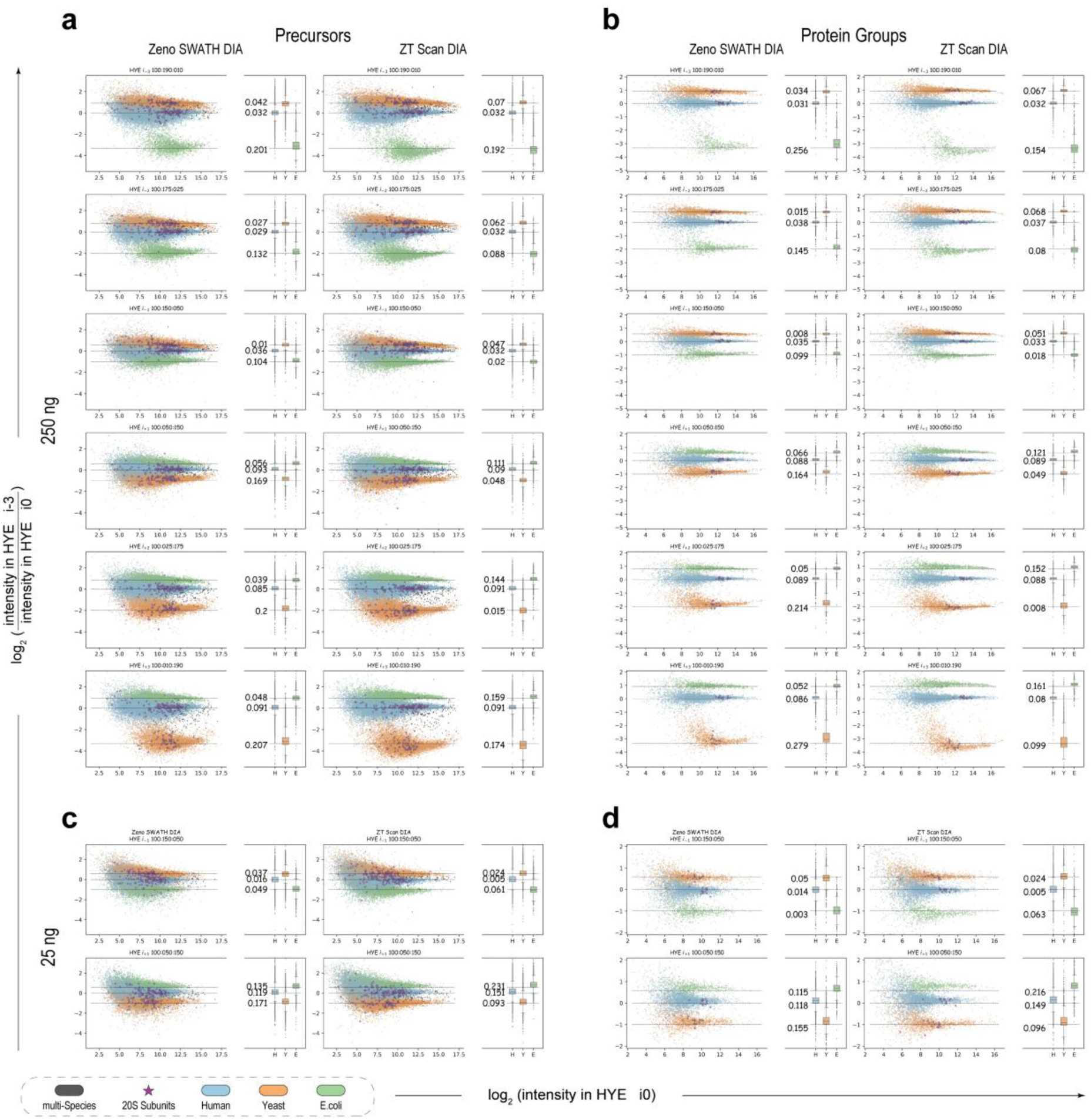
**ZT Scan DIA in a nanoflow separation configuration consistently offers greater quantitative accuracy,** demonstrated for **a**) precursors and **b**) protein groups. Log-ratio plots illustrate precursors and protein groups identified from 250 ng on-column loads. Panels **c** and **d** show the log-ratio plots of **c**) precursors and **d**) protein groups of 25 ng on-column loads. Note: Magenta stars in log-ratio plots depict 20S proteasome subunit precursors and proteins.

#### Empirical FDR estimates demonstrate robust peptide identification by both Zeno SWATH and ZT Scan DIA

We calculated experimental FDR values (see methods and Fig. 6a) in the LFQBench samples by searching human-only samples against the full multispecies spectral library. The precursor FDR values of both DIA methods remained between 0.44% to 0.62% without MBR, and is improved (i.e., decreased) to 0.37% to 0.59% with the implementation of MBR (hatched bars in Fig. 6b). Similarly, protein FDRs of both methods are controlled between 2.43% to 3.41% without MBR, and decreased to 2.21% to 2.71% with MBR implementation (Fig. 6c). With the improvements offered through the choice of nanoflow chromatographic separation, DIA-NN was able to implement MBR to not only increase similarity across replicates, but also, in most cases, to reduce the rate of falsely discovered precursors and proteins (hence, the downwards-facing hatched bars in Fig. 6c)

**Fig 6.**
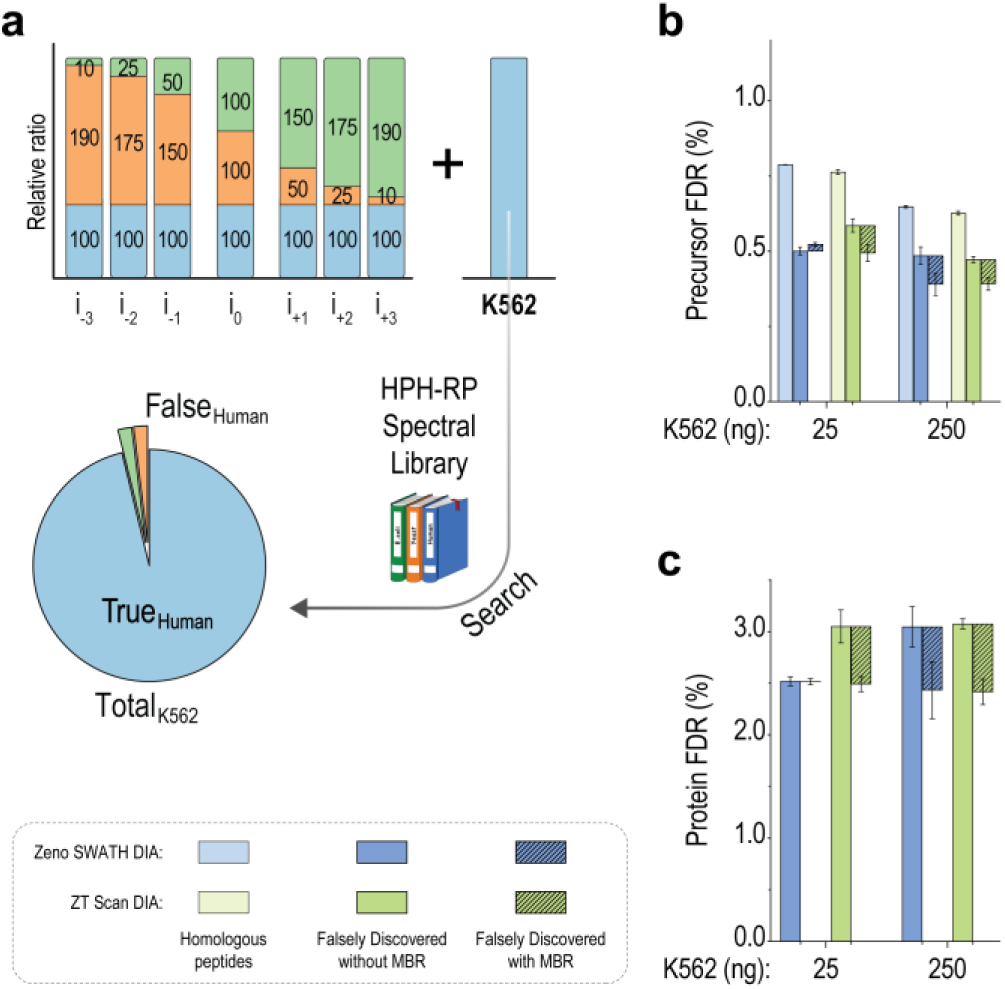
Empirical calculation of false discovery rates (FDR). **a**) The multispecies LFQBench design, particularly when accompanied by a species-assorted deep library, enables tracing all the identified precursors back to their species of origin, not only to calculate the ratio of falsely discovered precursors and proteins but also to distinguish homologous proteins with less ambiguity. The precursor and protein group FDR values of high-efficiency nano-flow (**b** & **c**) separation runs, calculated based on non-Human entries identified in a K562 (Human-only) sample, searched together with the LFQBench samples. A number of the flagged precursors were traced back to Human, as well as Yeast and/or E.coli in the HPH-RP library; these precursors were indicated as “homologous peptides” and since their true origin cannot be irrefutably ruled out, they were plotted separately (**b**). Note: Bar charts represent the mean of the measured values ± standard error of the mean (SEM) as error bars. The hatched bars represent searches where the match between run (MBR) is activated. These values (searches with MBR) originate from the mean of the corresponding searches without MBR. In other words, in cases where hatched bars are downward-facing (such as precursors identified in 250 ng K562 using ZT Scan DIA in panel (**b**)), activating MBR decreased the average of FDR values. (Note for formatting: The figure best suits a width of ∼8.8 cm)

#### Quantitative immunopeptidome analysis of pancreatic cancer using ZT Scan DIA

We recently demonstrated (25) the utility of Zeno SWATH DIA acquisition for tumor-associated-antigen target discovery in pancreatic cancer; we processed biological replicates of small pellets of 1M, 5M, and 10M Panc1 cells (cell equivalent on-column loads of approximately 160K, 800K, and 1.6M cells) and compared the benefits of DDA, as well as library-based DIA immunopeptidomic analyses. Here, we isolated HLA-peptides from a large pellet (500M cells) of Panc1 and injected three technical replicates of on-column loads equivalent to HLA-bound peptides isolated from 2.5M or 25M cells. Peptides were separated over a 15-min gradient (1% to 34% mobile phase B), at a flow rate of 150 nL/min, on an Aurora ELITE (15 cm x 75µm) XS UHPLC column (IonOpticks). Fig. 7 demonstrates the precursor (7- to 16-mer) and peptide (8- to 14-mer) counts (Fig. 7a) and identification reproducibility, i.e., data completeness and CV% values (Fig. 7b,c), all of which are comparable between using ZT Scan DIA and Zeno SWATH DIA. ZT Scan DIA identified 8157 to 8265 peptides in 2.5M and 11446 to 11697 peptides in 25M *cell equivalent* loads. ZenoSWATH DIA detected 7663 to 8143 peptides in 2.5M, and 11438 to 11591 peptides in 25M cell equivalent input injections. In other words, while Zeno SWATH DIA performs well, it was outperformed by ZT Scan DIA in smaller input samples, particularly in terms of quantitative reproducibility, i.e., CV% (Fig. 7b). Peptides identified with either of the methods show no deviations from the anticipated length distribution typical of HLA class I bound peptides (Fig. 7d), and exhibit striking similarity in Gibbs-cluster 9-mer core binding motifs (Fig. 7e), matching to HLA-allotypes expressed in Panc1.

**Fig 7.**
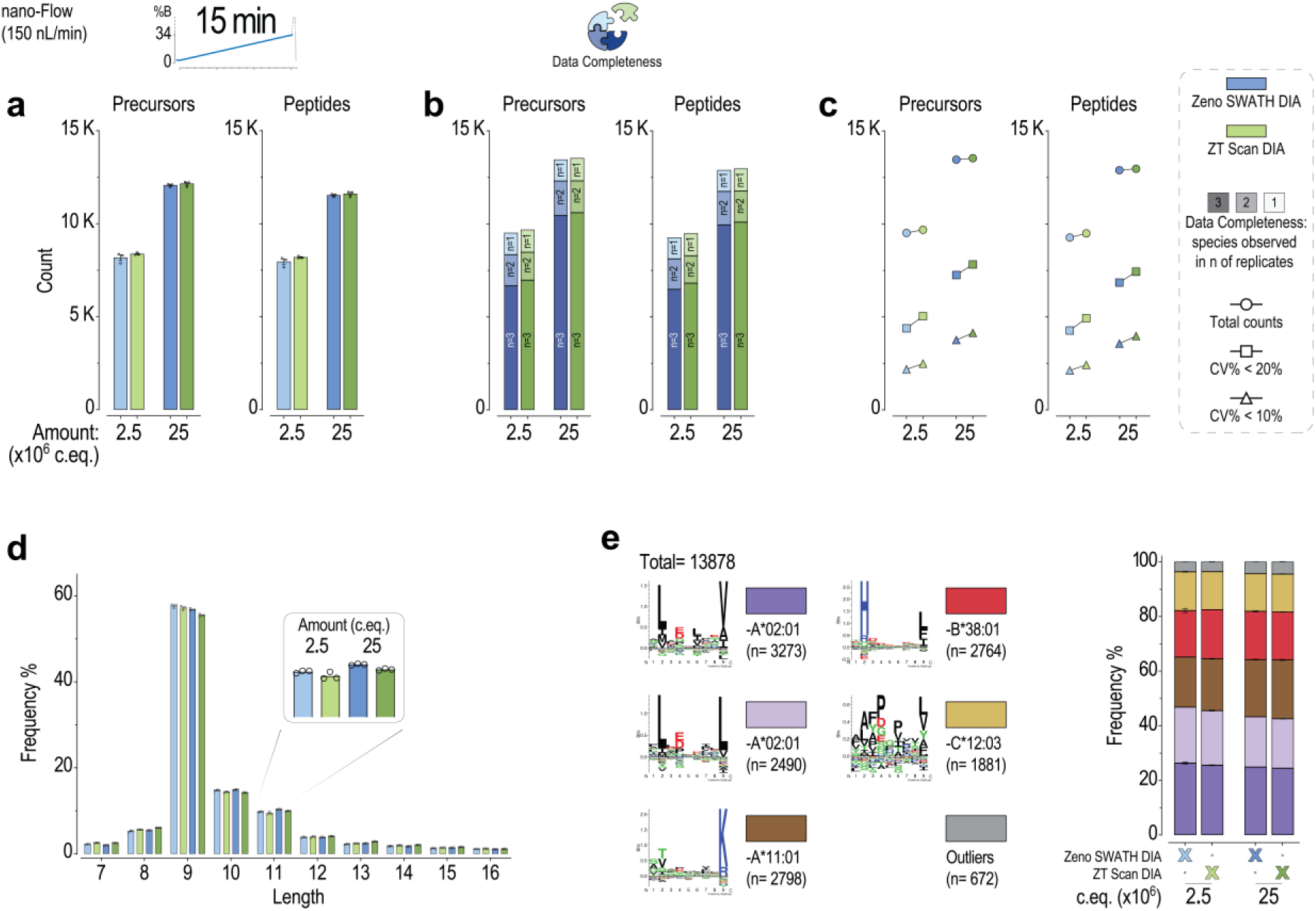
Analysis of Panc1 HLA peptidome. **a**) ZT Scan DIA and Zeno SWATH DIA offer a comparable identification depth at precursor and peptide (8- to 14-mers) levels. **b**) Both ZT Scan DIA and Zeno SWATH DIA provide reliable and reproducible identifications at the tested input amounts based on detection in the three replicates. **c**) ZT Scan DIA offers at least a modest improvement in the quantitative precision, at both precursor and peptide levels. **d**) HLA peptides identified in Panc1 across the four groups demonstrate predominant 9-mer-HLA-class I length distribution. **e**) Peptide sequences were clustered using the GibbsCluster 2.0 web tool. The resulting sequence motifs of peptides identified using both methods conform to the four HLA allotypes expressed in Panc1 (HLA-A*02:01, HLA-A*11:01, HLA-B*38:01, and HLA-C*12:03). Note: “Precursors” include different charge-states of 7- to 16-mer modified sequences after removing contaminants. “Peptides” are unique stripped sequences filtered for HLA class I length restriction (8- to 14-mers). The reason for including both was that the former has an analytical and technical importance, and the latter has biological relevance.

#### Identification and quantification are enhanced through the improved fragment-to-precursor assignment via the Q1 dimension of ZT scan DIA

FLDPRPLTV is a peptide derived from Cytochrome P450 1B1 (CYP1B1_190-198_) and is a known T cell epitope (42) observed in our data. In our previous study (25), we detected this peptide even in very small pellets (1M cell pellets = approximately 160K *cell equivalent* on-column) of Panc1. Fig. 8a,b show that during LC-MS/MS analysis, this epitope (FLDPRPLTV) co-elutes and co-fragments with KVFDPVPVGV, a peptide derived from ATP-dependent RNA Helicase A (DHX9_697-706_). The doubly charged monoisotopic precursors of these two peptides are 0.4977 m/z apart. In other words, the monoisotopic peak of FLDPRPLTV^++^ is only 7 ppm away from the first isotopomer of KVFDPVPVGV^++^ ([M - ^12^C + ^13^C + 2H]^++^), which guarantees their co-fragmentation in almost any condition (profile MS and centroided MS/MS scans in Fig. 8a,b). One of the values of DIA-MS in HLA-peptidomics is the ability to distinguish co-eluting and co-fragmenting precursors (with indiscernible precursor m/z), via fragment correlation over the elution profile of the peptides (25). Both ZT Scan and ZenoSWATH DIA perform equally well with this regard (XICs in Fig. 8a,b). Meanwhile, ZT Scan offers an additional dimension to distinguish the two precursors (Fig. 6c-e).

**Fig 8.**
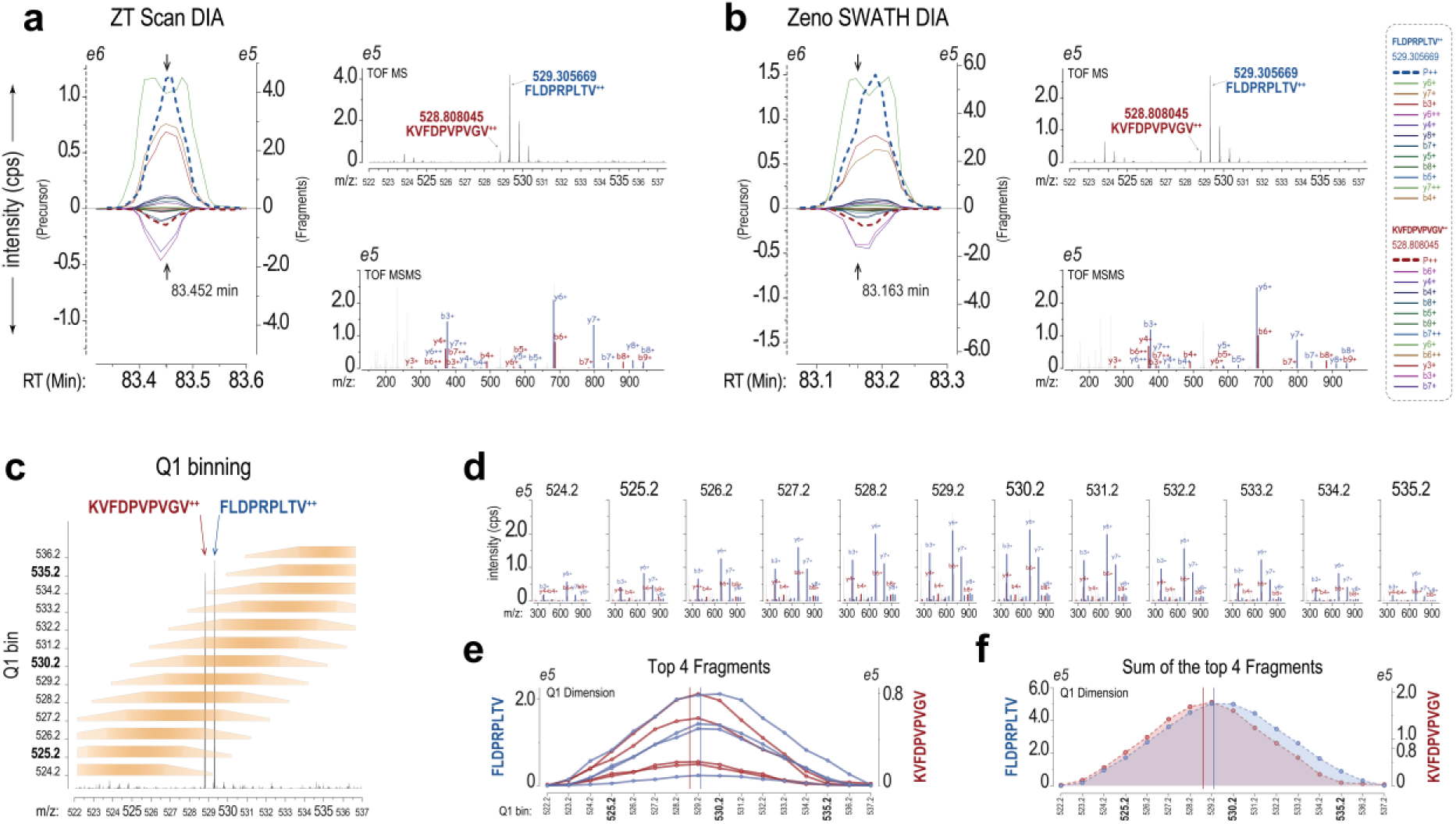
The utility of the Q1 dimension. **a** & **b**) The precursor and fragment extracted ion chromatograms (XIC), survey scans (profile spectra, top), and product scans (centroided spectra, bottom) of two peptides with potential for ambiguities in identification are illustrated. In the XIC plots, the left y-axes (dashed axes) show the precursor (dashed curves) intensities and the right y-axes (solid axes) depict fragment (solid lines) abundances; all y-axes are scaled to the most abundant precursor or fragment. Mass spectra depict precursor (top) and fragment (bottom) ions of FLDPRPLTV (blue) and KVFDPVPVGV (maroon). **c**-**f**) The scanning nature of the ZT Scan DIA offers an additional dimension to the LC-MS/MS data, the Q1 dimension, which, in the case of these two peptides, enhances the fragment-to-precursor assignment and enables the software to distinguish between the potentially ambiguous precursors, better quantification, as well as resolution of signals of interfering fragments. **c**) The diagram on the left shows the monoisotopic precursors of the two peptides (x-axis = precursor m/z) and the Q1 bins (scanning 5 m/z isolation windows collated into 1 m/z bins, shown in the y-axis). The full 5 m/z width of the isolation window is shown with uniform breadth. Since the isolation windows have non-square edges, the width of the illustrated isolation windows is extended by reducing breadths, accounting for the increasingly reduced ion transmissions in the borders. Note that other signals along the x-axis (including the isotopomers) are artificially removed to simplify the illustration. **d**) The sequential MS/MS spectra, traversing from left to right, belong to the corresponding Q1 bins in the left-side scheme. **e**) The emerging Q1 dimension is visualized for the top 4 fragments of each peptide, and **f**) the sum of the top 4 fragments. Left and right y-axes of the Q1 dimensions display the (summed-)intensities of FLDPRPLTV (blue) and KVFDPVPVGV (maroon), respectively.

In a single scan cycle of ZT Scan DIA, after acquiring the survey scan (TOF MS) for 50 ms, a 5 m/z isolation window smoothly slides up the m/z range at a rate of 0.75 m/z per millisecond, which is defined by the overall cycle time. Precursor ions are transmitted in accordance with the position of the Q1 within the mass range. The Q1 essentially moves independently of spectra being recorded with the acquisition engine signalling the start of each quadrupole cycle. Spectra of fragmented ions are continuously recorded and synchronized with the start of the quadrupole cycle (i.e., the portion of the cycle time spent on scanning the quadrupole for MS/MS acquisition). The quadrupole position for a given recorded spectrum is calibrated from residual parent ions in the fragmentation spectra (6). Over the elution profile of the two above-mentioned peptides (Fig. 8), at each scan cycle, the quadrupole isolation window first “meets” KVFDPVPVGV^++^, and then, after less than a millisecond, the right edge of the isolation window reaches FLDPRPLTV^++^. Since the isolation window in a quadrupole filter is not rectangular (16), the two precursors are transmitted slightly differently as they are being co-scanned. Fig. 8c shows a schematic diagram demonstrating how the Q1 window engages with and isolates the two precursors. The sequential MS/MS scans in Fig. 8d demonstrate how the intensity of each peptide’s fragments rise and fall over the Q1 bins between ∼524 and ∼535 m/z. When the top four fragments of each peptide are plotted against the Q1 bins (Fig. 8e), the “Q1 dimension” emerges and again distinguishes the two precursors. The distinction of the two precursors is even more observable in the Q1 dimension when these top four fragments are summed (Fig. 8f), allowing distinction of the signals for the two species at both MS1 and MS2 levels and improving quantitative accuracy.

## Discussion

For more than a decade, the rapidly evolving field of MS-based proteomics has benefited from several new technologies, in particular DIA-MS, that have advanced quantitative proteomics and peptidomics. The specificity and selectivity in a discrete-window DIA are achieved via narrower isolation windows to minimize signal interference from co-isolated analytes. However, the narrower the isolation window width is, the briefer the ion transmission becomes (43). In other words, to maximize specificity in a discrete-window DIA, one may inadvertently sacrifice assay sensitivity. Thus, there is a necessary compromise to find a “sweet spot” of isolation parameters in a discrete-window DIA assay that can achieve an optimum specificity without compromising sensitivity. Different approaches have been employed on different platforms to maximize both sensitivity and specificity. In Orbitraps, in both DDA and DIA modes, precursors are isolated with wider windows to maximize the sensitivity, while the higher specificity is often achieved due to higher MS1 precursor resolution. An example is the acquisition of wide-window DDA (WWA DDA) data with the intention to maximize the chimeric spectra, followed by a post hoc deconvolution with Chimerys (3) or MSFragger (4). On timsTOF platforms, even in DDA mode, precursors are isolated with windows of 2 to 3 m/z, while the 4^th^ dimension (ion mobility) can further disambiguate the chimeric spectra (4). It is worth mentioning that Synchro-PASEF (44) and midia-PASEF (45) are DIA acquisition modes used on timsTOF-based instruments, which exploit the full ion mobility dimension by scanning the quadrupole isolation window diagonally across the defined precursor m/z-ion mobility space (10). Similarly, in ZT Scan DIA, a 5 m/z isolation window scanned at 750 m/z per second is wide enough to maintain efficient ion transmission, while the added Q1 dimension enables fragment ions to be assigned to their corresponding precursors within effective theoretical windows of approximately 1 m/z. The increasing prevalence of scanning-quadrupole-enabled acquisition modes further underscores the growing need for search software to efficiently mine through the added dimensionality of the information-rich data for improved spectral deconvolution.

Here, we evaluated the performance of ZT Scan DIA in bottom-up proteomics, as well as immunopeptide identification and quantification. We believe that different DIA workflows should be compared on the same LC-MS platform so that differences in identification depth, quantitation, and peak sampling can be attributed to the acquisition method rather than to variation in chromatography, ionization efficiency, instrument sensitivity, resolving power, or scan speed. Thus, by employing a multi-species LFQBench approach (24), we observed improved quantitation accuracy with ZT Scan DIA acquisitions when compared to equivalent analyses using a more traditional sequential window approach on a ZenoTOF 7600+ system. These improvements were apparent both when focusing on microflow-based rapid chromatographic separations, as well as when we aimed for the more sensitive and higher chromatographic resolution obtained using nanoflow methods. We further assessed quantitative accuracy in a more challenging test by evaluating the log-ratios of protein complex subunit stoichiometry (46). Subunits of complexes such as the constitutive 20S proteasome are expected to be expressed at stoichiometric proportions (46). We demonstrated that ZT Scan DIA consistently measured the precursor and protein levels of 20S proteasome α and β subunits at ratios closer to their expected values. Additionally, across a wide range of parameters, ZT Scan DIA achieved tighter CV% distributions, demonstrating its superior quantitative reproducibility and precision compared to an equivalent variable window ZenoSWATH approach. This is in alignment with what Messner et al. (22) observed using Scanning-SWATH on a TripleTOF 6600 platform. Furthermore, we interrogated the LFQBench experiments using a purpose-made multi-species spectral library and demonstrated remarkably low experimentally measured precursor-level FDRs, often below 0.5%. We further investigated whether ZT Scan DIA was fit-for-purpose for HLA-peptide characterization. In line with our previous observations in comparing DDA and discrete-window DIA (25), our analyses highlighted that ZT Scan DIA and Zeno SWATH DIA provide an on-par ability to interrogate fragment XIC correlation. Yet, the scanning quadrupole (Q1) dimension of ZT scan DIA facilitates fragment-to-precursor assignment for further disambiguation of closely isolating immunopeptides. Given that the precursor m/z range that was sampled during the ZT Scan DIA method in our experiments was firmware-limited to 400 to 900 m/z, we expected that a subpopulation of singly charged precursors (900 to 1250 m/z) would be lost in these analyses.

Biological mixtures like cellular proteomes and immunopeptidomes encompass a wide dynamic range of analyte concentrations, with estimated numbers for proteins from hundreds to millions of copies, and immunopeptides from tens to a few hundred thousand copies per cell (47). To fathom this breadth of analytes with MS-based proteomics, regardless of how well the mass spectrometer acquires the data, a more efficient separation step is needed. It has long been demonstrated that, particularly using columns with narrower inner diameters and applying lower flow rates, improves chromatographic resolution of analytes, enhances ionization efficiency and MS response, thus resulting in a more sensitive quantitative analysis (41). Similarly, we observed that using a high-efficiency 60 cm separation column (Aurora Frontier XS, IonOpticks), not only increased the depth of identification (3-4 times) in low-input samples (1 ng of K562 digest), but also greatly improved the quantitative accuracy of both Zeno SWATH DIA and ZT scan DIA approaches. With choices of better chromatographic conditions, such as longer rather than shorter gradients, we detected on average 1.5 to 2.5 times more peptides per protein (expressed as % of sequence coverage).

The repertoire of immunopeptides presented in a given cell changes dynamically in response to external stimuli, such as cytokines, or disease conditions, including viral infections or during malignant transformations (48). For a T cell to mount an effective immune response against the target cell, the immunogenic epitope needs to be displayed on the cell surface and surpass a threshold, often reported as copies/cell (47–49). Thus, the differential quantity of immunopeptides in disease vs. healthy states may be used as one criterion to efficiently select targets. The bespoke immunopeptide ideally is not expressed on healthy tissues. However, a subthreshold expression on healthy cells may also be tolerated by T cells or TCR-mimicking therapeutics (47, 49, 50). Stopfer et al. (47) reviewed techniques, particularly labelling-based strategies, that have been adopted in the quantitative analysis of immunopeptidome, and critically discussed their challenges and limitations. Label-free quantification of immunopeptides using traditional DDA methods is often poorly reproducible, which cannot be mitigated using a match-between-run strategy due to the frequent presence of (near-)isobaric analytes and the potential for false identifications (25). Additionally, while DIA-MS is perceived to complicate the data analysis, it has several advantages that can empower immunopeptidome-derived target discovery. Label-free DIA-MS can improve reproducibility, has a superior sensitivity, and is less prone to issues such as ratio compression and expansion in a wider dynamic range, compared to label-based techniques (47, 51). Over the past couple of years, considerable effort has been devoted to evaluating the application of DIA-MS in immunopeptidome identification and quantification. Pak et al. (52) successfully demonstrated reproducible identification and quantification of immunopeptides using a variable-window DIA-MS on a Q-Exactive platform, with a striking 0.97 Pearson Correlation Coefficient across technical replicates. However, a challenge in quantitative immunopeptidomics is the variability introduced during immunoprecipitation and sample preparation, which can be evaluated and mitigated via MHC-bound heavy peptides as internal standards (47, 51, 53). Furthermore, as discussed above, a key aspect of the workflow is the software to search and interrogate the DIA data. Future studies are called for to evaluate the most recent advancements in DIA-MS for the quantification of immunopeptides, perhaps by utilizing MHC-bound heavy peptides (51) or approaches similar to that of Shabazy et al. (54).

The enhanced selectivity in data acquisition and the auxiliary effect of the additional Q1 dimension in ZT Scan DIA improve confidence in matching MS/MS data to peptide query parameters, whether using an experimental or an *in silico* predicted spectral library. The extended confidence in fragment-to-precursor assignment is reflected in improved fragment prioritization and lower false discoveries, which, in turn, result in a more accurate quantification. Several other parameters, including the dynamic range of analytes in the sample, input material amount, precursor m/z distribution, and chromatography-driven parameters like the average elution time calculated as the average of peak FWHM, can affect the choice of ZT Scan DIA parameters. Thus, further optimization for different types of samples is necessary to reach the best results. When taking a peptide-centric view of the data, as is often the case for quantitative immunopeptidomics, the additional Q1 dimension allowed unambiguous detection and quantitation of peptide species by resolving interfering signals arising from the co-isolation and co-fragmentation of the precursor ions.

## ASSOCIATED DATA

### Data availability

The mass spectrometry data for this project and the associated analyses have been deposited to the ProteomeXchange Consortium (http://proteomecentral.proteomexchange.org) via the PRIDE partner repository (55) with the dataset identifiers PXD069124, PXD069125, PXD069127, and PXD069132, respectively.

## Abbreviations

DIA: Data independent acquisition
MS: mass spectrometry
PASEF: parallel-accumulation sequential elution and fragmentation
SWATH-MS: sequential window acquisition of all theoretical mass spectra
LFQ: label-free quantitation

## Notes

### Competing Interest Statement

The authors declare the following financial interests/personal relationships which may be considered as potential competing interests: Anthony Purcell is a scientific advisor for Bioinformatics Solutions Inc (Canada), a shareholder and scientific advisor for Evaxion Biotech (Denmark), and a co-founder of Resseptor Therapeutics (Australia). The funders had no role in the design of the study; in the collection, analyses, or interpretation of data; in the writing of the manuscript; or in the decision to publish the results. I.B., J.C., P.P., K.T., and S.T. are employees of SCIEX. A.P. and J.J.S. are employees of IonOpticks. Both companies provided early access to technology that is now commercially available. Neither had influence over the decision to publish.

### Funding

- NHMRC: 2016596, 2013631, 2029227

