## Supplementary material for "Zeno Trap-enabled Quadrupole-Scanning Data-Independent Acquisition in nanoflow Quantitative Proteomics and Immunopeptidomics": Supp Info

***Table S1.*** *Theoretical amounts of each species' peptides in each test sample mixture based on the ratios.*

| | 25 ng | 50 ng | 250 ng |
|---|---|---|---|
| ***i±1:*** | | | |
| **100** | 8.33 ng | 16.67 ng | 83.33 ng |
| **150** | 12.5 ng | 25 ng | 125 ng |
| **50** | 4.17 ng | 8.33 ng | 41.67 ng |
| ***i±2:*** | | | |
| **100** | 8.33 ng | 16.67 ng | 83.33 ng |
| **175** | 14.58 ng | 29.17 ng | 145.83 ng |
| **25** | 2.08 ng | 4.17 ng | 20.83 ng |
| ***i±3:*** | | | |
| **100** | 8.33 ng | 16.67 ng | 83.33 ng |
| **190** | 15.83 ng | 31.67 ng | 158.33 ng |
| **10** | 0.83 ng | 1.67 ng | 8.33 ng |

***Table S2.*** *Summary of chromatographic conditions and parameters.*

| *Experiment* | *Total run (min)* | *Active Gradient (min)* | *Active Gradient (B%)* | *Flow rate (nL/min)* | *UHPLC Column* |
|---|---|---|---|---|---|
| Serial Dilution and LFQBench with micro-flow separations | 11 | 1 | 3 to 30 | 5000 | 150 mm x 300 µm kinetex (Phenomenex) |
| | 13 | 3 | 3 to 30 | 5000 | 150 mm x 300 µm kinetex (Phenomenex) |
| | 15 | 5 | 3 to 30 | 5000 | 150 mm x 300 µm kinetex (Phenomenex) |
| | 27 | 15 | 3 to 30 | 5000 | 150 mm x 300 µm kinetex (Phenomenex) |
| Serial Dilution and LFQBench with nano-flow separations | 115 | 30 | 1 to 34 | 150 | 600 mm x 75 µm Aurora FRONTIER XS (IonOpticks) |
| HPH-RP Library with nano-flow | 185 | 92 | 1 to 34 | 150 | 600 mm x 75 µm Aurora FRONTIER XS (IonOpticks) |
| HLA-peptide analysis with nano-flow separations | 100 | 15 | 1 to 34 | 150 | 150 mm x 75 µm Aurora ELITE (IonOpticks) |

***Table S3.*** *Summary of MS/MS acquisition methods' parameters (Dyn. excl. = dynamic exclusion, occur. = occurrence).*

| ***Experiment*** | ***Acquisition mode*** | ***TOF spectra ranges*** | ***Specific parameters*** | ***Dwell and cycle times*** |
|---|---|---|---|---|
| Assessment of tryptic peptides quantitative precision and accuracy | ZT Scan DIA | MS = 400-900 mz<br>MS/MS = 140-1750 mz | Sliding window = 5 mz<br>Q1 Scanning Speed = 750 mz/s | Dwell = 6.7 ms<br>Total cycle time = 782 ms |
| | | | Sliding window = 10 mz<br>Q1 Scanning Speed = 750 mz/s | Dwell = 13.3 ms<br>Total cycle time = 789 ms |
| | ZenoSWATH DIA | MS = 400-900 mz<br>MS/MS = 140-1800 mz | Number of windows = 60<br>Window size = 9-10 mz | Accumulation time = 7 ms<br>Total cycle time = 786 ms |
| | | | Number of windows = 60<br>Window size = 9-10 mz | Accumulation time = 13 ms<br>Total cycle time = 1146 ms |
| HYE HPH-RP Spectral Library generation HLA-peptide analysis | DDA | MS = 400-1500 mz<br>MS/MS = 140-1800 mz | Top N = 45 most abundant<br>Dyn. excl. = 5 s / 1 occur.<br>Charge = +2 to +5 | Accumulation time = 20 ms<br>Total cycle time = 1259 ms |
| | ZT Scan DIA | MS = 400-900 mz<br>MS/MS = 140-1750 mz | Sliding window = 5 mz<br>Q1 Scanning Speed = 750 mz/s | Dwell = 6.7 ms<br>Total cycle time = 782 ms |
| | ZenoSWATH DIA | MS = 400-900 mz<br>MS/MS = 140-1800 mz | Number of windows = 65<br>Minimum window size = 5 mz | Accumulation time = 7 ms<br>Total cycle time = 843 ms |

***Table S4.*** *Isolation scheme of the ZenoSWATH method designed to scan between 400 m/z to 900 m/z.*

| Window start | Window stop | Window start | Window stop |
|---|---|---|---|
| 399.5 | 411 | 566.5 | 571.5 |
| 410 | 421 | 570.5 | 575.5 |
| 420 | 430.5 | 574.5 | 579.5 |
| 429.5 | 439.5 | 578.5 | 583.5 |
| 438.5 | 447 | 582.5 | 587.5 |
| 446 | 454 | 586.5 | 591.5 |
| 453 | 459.5 | 590.5 | 595.5 |
| 458.5 | 465 | 594.5 | 599.5 |
| 464 | 470 | 598.5 | 604 |
| 469 | 474.5 | 603 | 608.5 |
| 473.5 | 479 | 607.5 | 613 |
| 478 | 483.5 | 612 | 618 |
| 482.5 | 487.5 | 617 | 623.5 |
| 486.5 | 491.5 | 622.5 | 629 |
| 490.5 | 495.5 | 628 | 635 |
| 494.5 | 499.5 | 634 | 641 |
| 498.5 | 503.5 | 640 | 647 |
| 502.5 | 507.5 | 646 | 654 |
| 506.5 | 511.5 | 653 | 661 |
| 510.5 | 515.5 | 660 | 669 |
| 514.5 | 519.5 | 668 | 679 |
| 518.5 | 523.5 | 678 | 690 |
| 522.5 | 527.5 | 689 | 703 |
| 526.5 | 531.5 | 702 | 720 |
| 530.5 | 535.5 | 719 | 740 |
| 534.5 | 539.5 | 739 | 763 |
| 538.5 | 543.5 | 762 | 788 |
| 542.5 | 547.5 | 787 | 813 |
| 546.5 | 551.5 | 812 | 837 |
| 550.5 | 555.5 | 836 | 860 |
| 554.5 | 559.5 | 859 | 881.5 |
| 558.5 | 563.5 | 880.5 | 900.5 |
| 562.5 | 567.5 | 566.5 | 571.5 |
| 399.5 | 411 | 570.5 | 575.5 |

***Table S5.*** *Spectral library generation parameters in FragPipe 22.0.*

| ***Parameter*** | ***value*** |
|---|---|
| MSFragger (common options): | |
| Precursor mass tolerance | 20 ppm |
| Fragment mass tolerance | 20 ppm |
| Isotope error | 0/1/2/3 |
| Enzyme | Stricttrypsin |
| Clip N-term M | Yes |
| Peptide length | 7 to 50 |
| Peptide mass (mass) | 500 to 5000 |
| MSFragger (modifications): | |
| Max variable modification | 3 |
| Variable modification 1 | M Oxidation= 15.9949 (max. 3 per peptide) |
| Variable modification 2 | Protein N-terminus acetylation= 42.0106 (max. 1 per peptide) |
| Fixed modification | C-Carbamidomethylation= 57.02146 |
| Validation (MSBooster): | |
| Predict RT | DIANN |
| Predict spectra | DIANN |
| Validation (PSM validation): | |
| Percolator | default |
| Validation (FDR fiter and report): | |
| Ion FDR (--ion) | 0.01 |
| PSM FDR (--psm) | 0.01 |
| Peptide FDR (--pept) | 0.01 |
| Protein FDR (--prot) | 0.01 |
| IonQuant (LFQ) tab | |
| Turned off | |
| Spectral Library tab | |
| RT Lowess Fraction | 0 |
| Max fragment tolerance | 15 ppm |
| Max UniMod tolerance | 0.02 Da |
| RT calibration | Automatic selection of a run as a reference RT |
| IM calibration | N/A |
| Fragment types | b- and y-ions |

***Table S6.*** *DIA-NN parameters to search Human (K562) tryptic peptide serial dilutions.*

| ***Parameter*** | ***value*** |
|---|---|
| Spectral Library and Database: | |
| Spectral Library | SCIEX K562+HeLa HPH-RP library |
| FASTA | nil |
| Reannotate | nil |
| Contaminants | nil |
| Precursor ion generation: | |
| FASTA digest … | nil |
| Deep learning … | nil |
| Protease | Trypsin/P |
| Missed cleavages | 1 |
| Max var. mod. | 0 |
| N-term Met excision | Yes |
| C carbamidomethylation | Yes |
| Oxidation of Met | nil |
| Other mods | nil |
| Peptide length range | 7 to 30 |
| Peptide charge range | +1 to +4 |
| Precursor m/z range | 400 to 900 |
| Fragment io m/z range | 140 to 1800 |
| Output: | |
| Reuse .quant files | nil |
| Quantities matrices | Yes (unique folders specified) |
| XICs | nil |
| Generate Spec. Lib. | Yes or nil (not important) |
| Generate prosit input | nil |
| Precursor FDR | 1% |
| Algorithm: | |
| Mass (MS2) accuracy | 20 ppm |
| MS1 accuracy | 0 (automatic inference) |
| Scan window (=RT radius) | 6 |
| Unrelated runs | nil |
| Peptidoforms | nil |
| MBR | Yes |
| Heuristic protein inference | Yes |
| No shared spectra | Yes |
| Protein inference | Genes |
| Machine-learning | Single-pass |
| Quantification strategy | QuantUMS (high precision) |
| Cross-run normalization | RT-dependent |
| Library generation | Smart profiling |
| Speed and RAM | Optimal results |
| Additional Options: | |
| --report-lib-info | For both ZenoSWATH DIA and ZT Scan DIA files |
| --scanning-swath | For ZT Scan DIA files |

***Table S7.*** *DIA-NN 1.9.2 parameters to search LFQBench data in a library-based mode.*

| *Parameter* | | *value* |
|---|---|---|
| **For *in silico* library generation:** | | |
| Spectral Library and Database: | | |
| | Spectral Library | HYE HPH-RP library (FragPipe output) |
| Precursor ion generation: | | |
| | Deep learning … | Yes |
| Output: | | |
| | Generate Spec. Lib. | Yes |
| **For main searches using the *in silico* library:** | | |
| Spectral Library and Database: | | |
| | Spectral Library | HYE HPH-RP DIANN in silico library (deep learning refined) |
| Precursor ion generation: | | |
| | Deep learning … | nil |
| Output: | | |
| | Generate Spec. Lib. | nil |
| **For both searches:** | | |
| | FASTA | nil |
| | Reannotate | nil |
| | Contaminants | nil |
| Precursor ion generation: | | |
| | FASTA digest … | nil |
| | Protease | Trypsin/P |
| | Missed cleavages | 1 |
| | Max var. mod. | 1 |
| | N-term Met excision | Yes |
| | C carbamidomethylation | Yes |
| | Oxidation of Met | Yes |
| | Other mods | nil |
| | Peptide length range | 7 to 30 |
| | Peptide charge range | +1 to +4 |
| | Precursor m/z range | 400 to 900 |
| | Fragment io m/z range | 140 to 1800 |
| Output: | | |
| | Reuse .quant files | nil |
| | Quantities matrices | Yes (unique folders specified) |
| | XICs | nil |
| | Generate prosit input | nil |
| | Precursor FDR | 1% |
| Algorithm: | | |
| | Mass (MS2) accuracy | 20 ppm |
| | MS1 accuracy | 0 (automatic inference) |
| | Scan window (=RT radius) | 6 |
| | Unrelated runs | nil |
| | Peptidoforms | nil |
| | MBR | Yes |
| | Heuristic protein inference | Yes |
| | No shared spectra | Yes |
| | Protein inference | Genes |
| | Machine-learning | Double-pass |
| | Quantification strategy | QuantUMS (high accuracy) |
| | Cross-run normalization | RT-dependent |
| | Library generation | Smart profiling |
| | Speed and RAM | Optimal results |
| Additional Options: | | |
| | --report-lib-info | For both ZenoSWATH DIA and ZT Scan DIA files |
| | --scanning-swath | For ZT Scan DIA files |

***Table S8.*** *DIA-NN 1.9.2 parameters to search Panc1 HLA-peptidome data in a library-based mode.*

| ***Parameter*** | ***value*** |
|---|---|
| **For *in silico* library generation:** | |
| Spectral Library and Database: | |
| Spectral Library | HLA-p HPH-RP library published in PXD054360 (FragPipe output) |
| Precursor ion generation: | |
| Deep learning … | Yes |
| Output: | |
| Generate Spec. Lib. | Yes |
| **For main searches using the *in silico* library:** | |
| Spectral Library and Database: | |
| Spectral Library | DIANN *in silico* library (deep learning refined) |
| Precursor ion generation: | |
| Deep learning … | nil |
| Output: | |
| Generate Spec. Lib. | nil |
| **For both searches:** | |
| FASTA | nil |
| Reannotate | nil |
| Contaminants | nil |
| Precursor ion generation: | |
| FASTA digest … | nil |
| Protease | Trypsin/P (Does not change the results) |
| Missed cleavages | 2 |
| Max var. mod. | 3 |
| N-term Met excision | Yes |
| C carbamidomethylation | Yes (Does not change the results) |
| Oxidation of Met | Yes |
| Other mods | nil |
| Peptide length range | 7 to 30 |
| Peptide charge range | +1 to +4 |
| Precursor m/z range | 300 to 1800 (DIANN will recognize and use the acquired range) |
| Fragment io m/z range | 100 to 1800 (DIANN will recognize and use the acquired range) |
| Output: | |
| Reuse .quant files | nil |
| Quantities matrices | Yes (unique folders specified) |
| XICs | nil |
| Generate prosit input | nil |
| Precursor FDR | 1% |
| Algorithm: | |
| Mass (MS2) accuracy | 20 ppm |
| MS1 accuracy | 20 ppm |
| Scan window (=RT radius) | 0 (automatic inference) |
| Unrelated runs | nil |
| Peptidoforms | nil |
| MBR | nil |
| Heuristic protein inference | Yes |
| No shared spectra | Yes |
| Protein inference | Off |
| Machine-learning | Double-pass |
| Quantification strategy | QuantUMS (high precision) |
| Cross-run normalization | RT-dependent |
| Library generation | Smart profiling |
| Speed and RAM | Optimal results |
| Additional Options: | |
| --report-lib-info | For both ZenoSWATH DIA and ZT Scan DIA files |
| --scanning-swath | For ZT Scan DIA files |
| --var-mod UniMod:7,0.984,NQ | For both ZenoSWATH DIA and ZT Scan DIA files |
| --var-mod UniMod:312,119.0041,C | For both ZenoSWATH DIA and ZT Scan DIA files |

***Table S9.*** *The list of Human and Yeast 20S Proteasome subunits, used for evaluating quantitative accuracy based on protein 20S Proteasome subunit stoichiometry. Note that the three underscored beta subunits PSMB5 (β5), PSMB6 (β1), and PSMB7 (β2) are the catalytic subunits of the constitutive 20S proteasome.*

| ***α-Subunits*** | ***Human*** | ***Yeast*** | ***β-Subunits*** | ***Human*** | ***Yeast*** |
|---|---|---|---|---|---|
| **PSMA1 (α6):** | P25786 | P21243 | **PSMB1 (β6):** | P20618 | P38624 |
| **PSMA2 (α2):** | P25787 | P23639 | **PSMB2 (β4):** | P49721 | P25043 |
| **PSMA3 (α7):** | P25788 | P23638 | **PSMB3 (β3):** | P49720 | P25451 |
| **PSMA4 (α3):** | P25789 | P40303 | **PSMB4 (β7):** | P28070 | P22141 |
| **PSMA5 (α5):** | P28066 | P32379 | **PSMB5 (β5):** | P28074 | P30656 |
| **PSMA6 (α1):** | P60900 | P40302 | **PSMB6 (β1):** | P28072 | P23724 |
| **PSMA7 (α4):** | O14818 | P21242 | **PSMB7 (β2):** | Q99436 | P30657 |

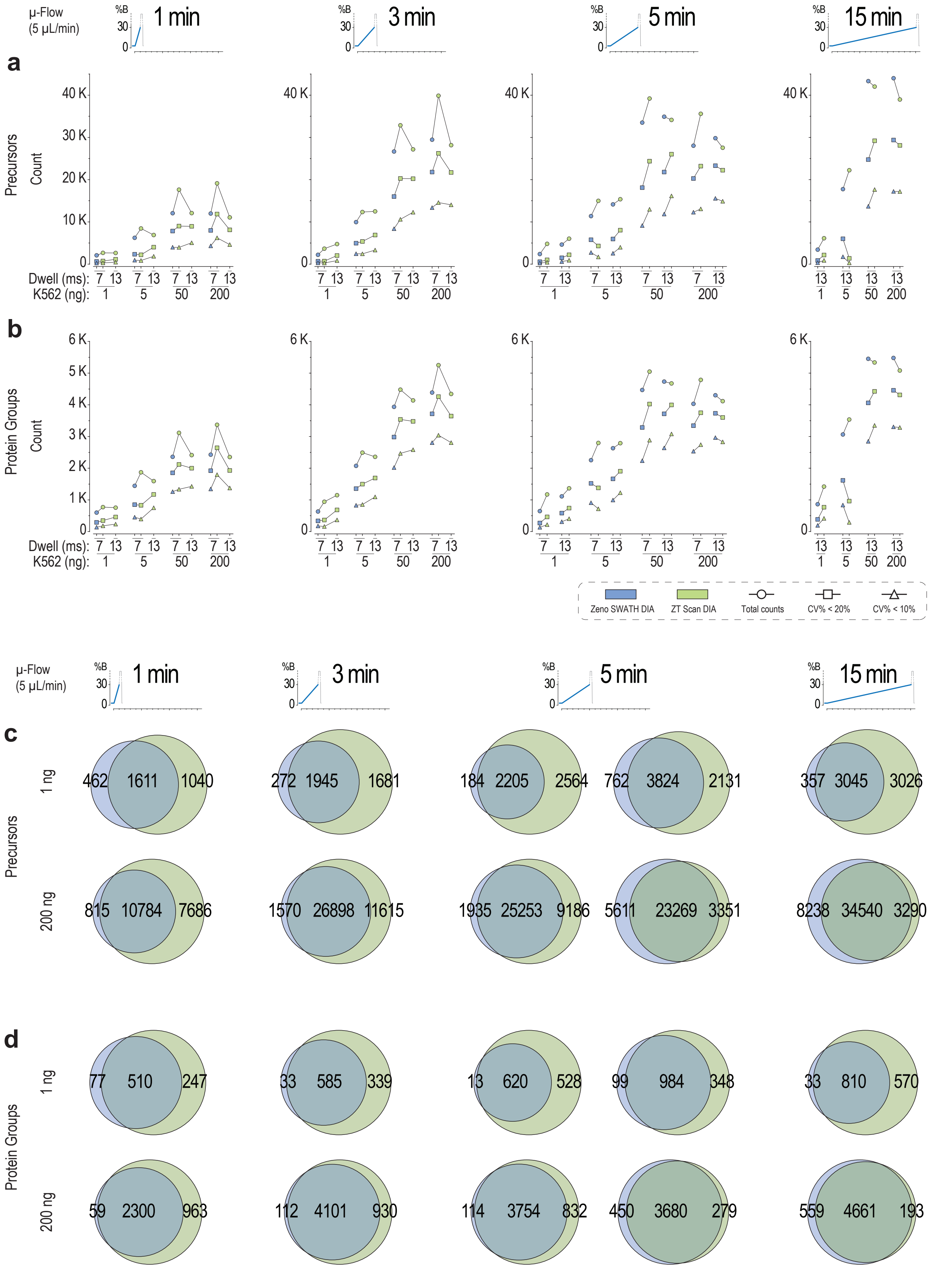

***Fig. S1.*** *ZT Scan DIA overall improves the depth of identification with better precision, at **a**) precursor **b**) protein group levels with lower input amounts (on-column loads) and shorter gradients (i.e., higher throughputs). Circles represent total protein groups and precursor counts of ZT Scan DIA (green) and Zeno SWATH DIA (blue) acquisitions of serially diluted (1, 5, 50, 200 ng) Human tryptic digest (K562 cell line). Squares and triangles represent counts of entries with a CV% less than 20% and 10%, respectively. Gradient lengths are shown by the keys on top of each protein-precursor plot pair (vertically); from left to right, 1, 3, 5, and 15 min separations at a 5 μL/min flow rate.*

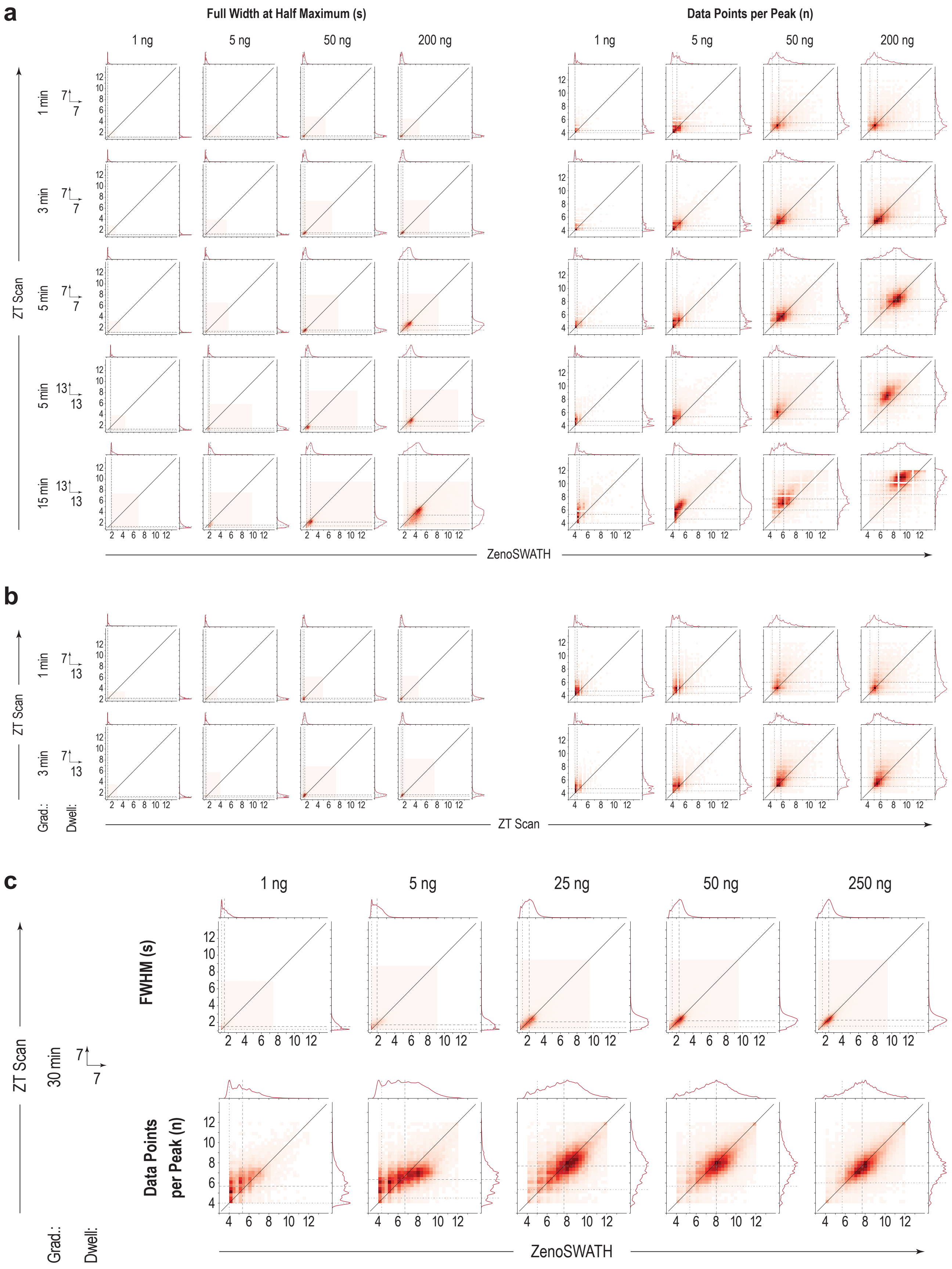

***Fig. S2*** *Summary of the two-by-two comparisons of Peak widths, expressed as the full-width at half maximum (FWHM) in the left set of plots for the microflow K562 experiments. The right set of plots illustrates the comparisons of the number of points across the peak (base-to-base). Both sets of values were calculated as the average of replicates for each precursor.* ***a****) This panel compares the ZenoSWATH DIA vs. the ZT Scan DIA with similar separation and acquisition parameters.* ***b****) This panel illustrates comparisons of the two different Dwell times of ZT Scan DIA, yet with the same gradient lengths and on-column loads.* ***c****) Summary of the two-by-two comparisons of Peak widths, expressed as the full-width at half maximum (FWHM) in the top set of plots for the nanoflow K562 experiments. The bottom set of plots illustrates the comparisons of the number of points across the peak (base-to-base). Both sets of values were calculated as the average of replicates for each precursor. Comparisons were made between ZenoSWATH DIA (x-axis) vs. the ZT Scan DIA (y-axis). Note: The on-column loads are written at the top of the figure. The gradient length (Grad.) is written in minutes on the left side of the figure. The dwell times of the method on the x-axis and the y-axis are illustrated with the two small perpendicular arrows next to the gradient length. In each plot, the histograms on the top and right axis of the plot show the distributions in the corresponding dimension. The x=y line depicts the ideal conditions where both methods would have performed exactly equally. Dashed lines also mark the 10th percentile and the median of the observed values for each method.*

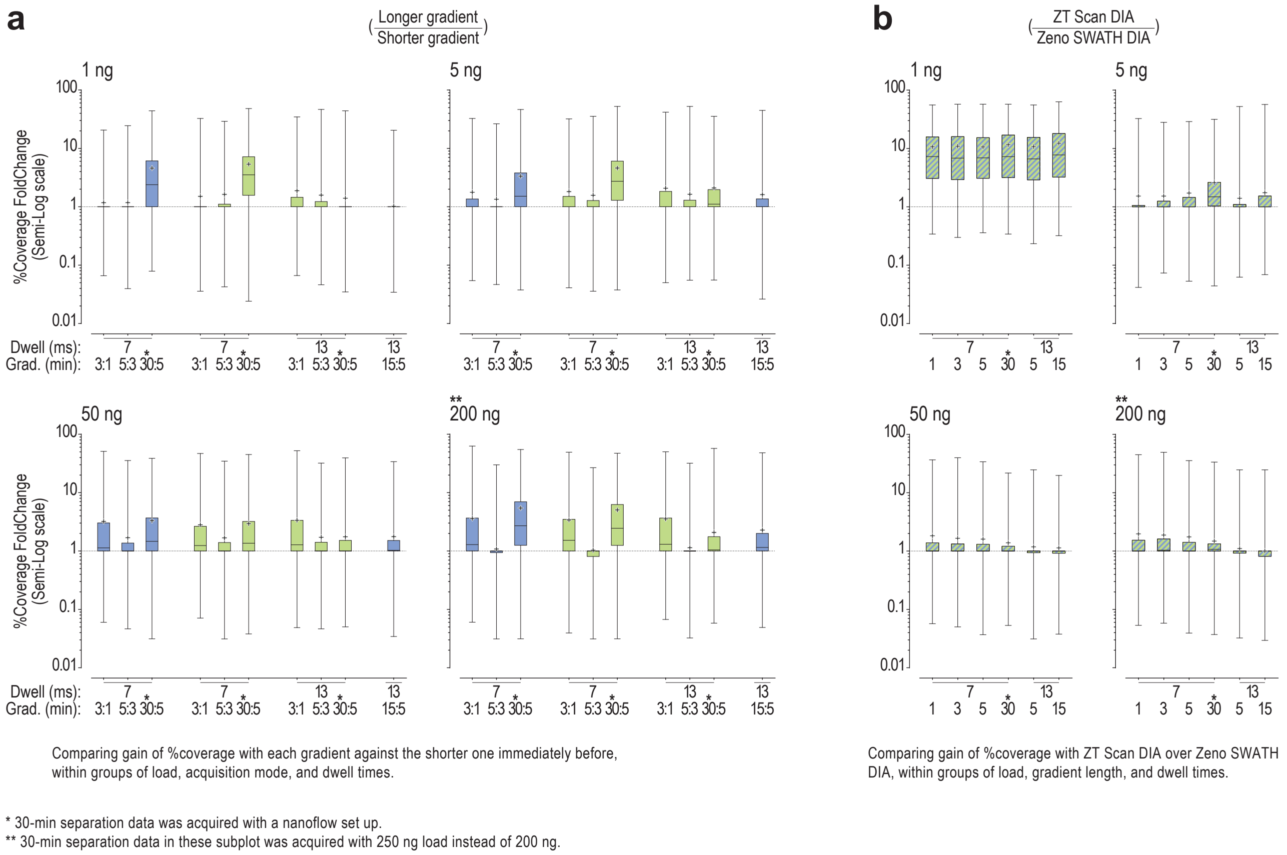

***Fig. S3.*** *Improved chromatographic conditions and the choice of ZT Scan DIA boost the protein sequence coverage.* ***a****) The K562 serial dilution results were grouped based on the on-column loads, acquisition methods, and dwell times. Fold changes of protein sequence coverage was calculated for the choice of each gradient length, over the shorter one immediately before; the expression "3:1", for instance, represents the fold change of 3-min over 1-min gradient.* ***c****) The K562 serial dilution results were grouped based on the on-column loads, gradient lengths, and dwell times, and the fold change of protein coverage was calculated for the choice of ZT Scan over ZenoSWATH as the acquisition mode. Each box plot spans the interquartile range, with a solid line at the median and the whiskers marking the minimum and maximum. Solid "+" marks demonstrate the arithmetic mean. Note: the data for 1, 3, 5, and 15 min gradients were acquired in a microflow setup, while the 30 min gradient was acquired in a nanoflow separation. Additionally, 30-min gradient separation results in the "200 ng" subplots were acquired with 250 ng loads instead of 200 ng.*

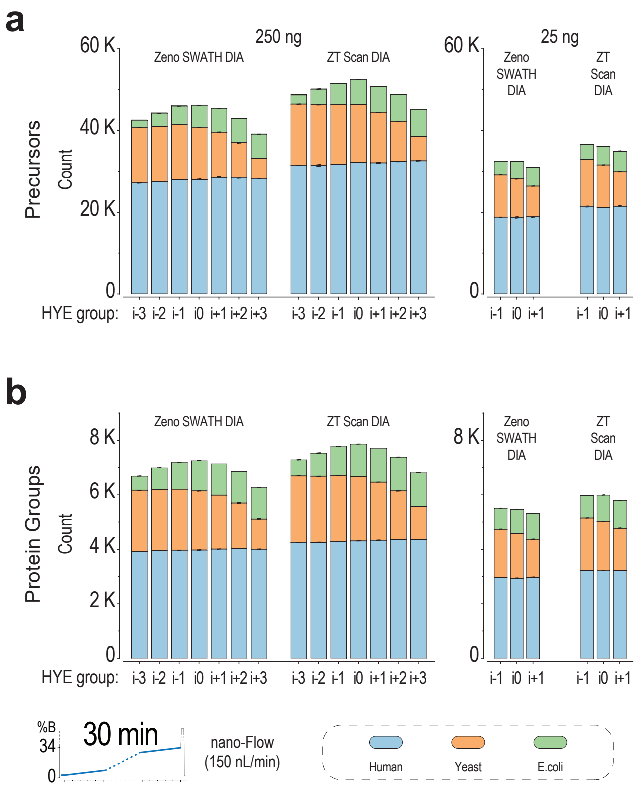

***Fig. S4.*** *The number of **a**) precursors and **b**) protein groups identified per species in each LFQBench group is shown. The on-column loads (25 or 250 ng) are written over the subplots. For each subplot, the acquisition modes are bannered over the left bar sets (Zeno SWATH DIA) and right bar sets (ZT Scan DIA). The dwell time for all acquisitions was 7 ms. (Note: Bar charts represent the mean of the measured values ± standard error of the mean (SEM) as error bars.)*
